# Human assembloid model of the ascending neural sensory pathway

**DOI:** 10.1101/2024.03.11.584539

**Authors:** Ji-il Kim, Kent Imaizumi, Mayuri Vijay Thete, Zuzana Hudacova, Ovidiu Jurjuţ, Neal D. Amin, Grégory Scherrer, Sergiu P. Paşca

**Affiliations:** Department of Psychiatry and Behavioral Sciences, Stanford University, Stanford, CA 94305, USA; Stanford Brain Organogenesis Program, Wu Tsai Neurosciences Institute & Bio-X, Stanford, CA 94305, USA; Department of Cell Biology and Physiology, Department of Pharmacology, UNC Neuroscience Center, The University of North Carolina, Chapel Hill, NC 27599, USA

**Author notes:** Correspondence to (S.P.P.). These authors contributed equally (Ji-il Kim, Kent Imaizumi).

## Abstract

The ascending somatosensory pathways convey crucial information about pain, touch, itch, and body part movement from peripheral organs to the central nervous system. Despite a significant need for effective therapeutics modulating pain and other somatosensory modalities, clinical translation remains challenging, which is likely related to species-specific features and the lack of in vitro models to directly probe and manipulate this polysynaptic pathway. Here, we established human ascending somatosensory assembloids (hASA)– a four-part assembloid completely generated from human pluripotent stem cells that integrates somatosensory, spinal, diencephalic, and cortical organoids to model the human ascending spinothalamic pathway. Transcriptomic profiling confirmed the presence of key cell types in this circuit. Rabies tracing and calcium imaging showed that sensory neurons connected with dorsal spinal cord projection neurons, which ascending axons further connected to thalamic neurons. Following noxious chemical stimulation, single neuron calcium imaging of intact hASA demonstrated coordinated response, while four-part concomitant extracellular recordings and calcium imaging revealed synchronized activity across the assembloid. Loss of the sodium channel SCN9A, which causes pain insensitivity in humans, disrupted synchrony across the four-part hASA. Taken together, these experiments demonstrate the ability to functionally assemble the essential components of the human sensory pathway. These findings could both accelerate our understanding of human sensory circuits and facilitate therapeutic development.

## Introduction

The ascending somatosensory pathways are responsible for conveying environmental and bodily sensory information from peripheral organs to the central nervous system. Somatosensory neurons, whose cell bodies reside in dorsal root ganglia (DRG) and trigeminal ganglia, are pseudo–unipolar cells that innervate peripheral organs including the skin and transmit somatosensory signals to the spinal cord. Dorsal spinal cord neurons monosynaptically transmit this information to various brain structures, including the thalamus, which further relays these signals to the cerebral cortex^1,2^. Genetic or environmental disruptions to these pathways can lead or contribute to various neurological disorders, including chronic pain and autism spectrum disorder, and impact both heightened and diminished sensory functions^3–6^. Despite the clinical significance, how ascending sensory pathways form during human development and how functional defects arise in pathological conditions are still poorly understood. Challenges in bench to bedside translation of pain and sensory research are thought to be related, at least in part, to differences across species. Crucially, the discovery of novel treatments and their translation is hindered by the lack of reliable human experimental models reconstituting the ascending sensory pathway^7^. In fact, concomitantly probing all components of these circuits via imaging or recording has not yet been achieved in animal models, which limits our understanding of the consequences of various sensory manipulations at the circuit level. Recent advances in organoid and assembloid technologies hold promise for generating self-organizing, stem cell-based models to study and manipulate human neural circuits *ex vivo* in the context of development as well as disease states. We previously developed three-part assembloids to recapitulate the descending cortico–spino–muscle pathway connecting deep layer cortical glutamatergic neurons to cholinergic spinal cord motor neurons to muscle^8^. Generation of functional circuits in assembloids with more than three components, which would enable the assembly of the ascending sensory pathway from sensory neurons to cerebral cortex, has not yet been achieved.

Here, we focus on the anterolateral spinothalamic pathway, which is particularly prominent in humans^9^ and is crucial for conveying pain and other sensory information through the spinal cord and thalamus^1^. In this circuit, sensory neurons in the dorsal root ganglia (DRG) connect with glutamatergic neurons in the dorsal spinal cord, which relay information to thalamus via monosynaptic connections and subsequently to the somatosensory cortex and other cortical areas (Fig. 1a). To reconstitute this pathway from its components, we separately generated somatosensory organoids, dorsal spinal cord organoids, diencephalic organoids, and cerebral cortical organoids from pluripotent stem cells and then functionally assembled them to create four-part human ascending somatosensory assembloids (hASA). We demonstrated that chemical stimulation or glutamate uncaging of sensory neurons elicited responses across within hASA. Calcium imaging and electrophysiological recordings of these assembloids revealed synchronized activity across the ascending pathway. Lastly, we applied this platform to study circuit–level consequences of loss of the Na_v_1.7 sodium channel encoded by the *SCN9A* gene, which has been associated with congenital insensitivity to pain and other neurological disorders, and discovered disrupted synchronized activity across the four-part hASA.

**Fig. 1.**
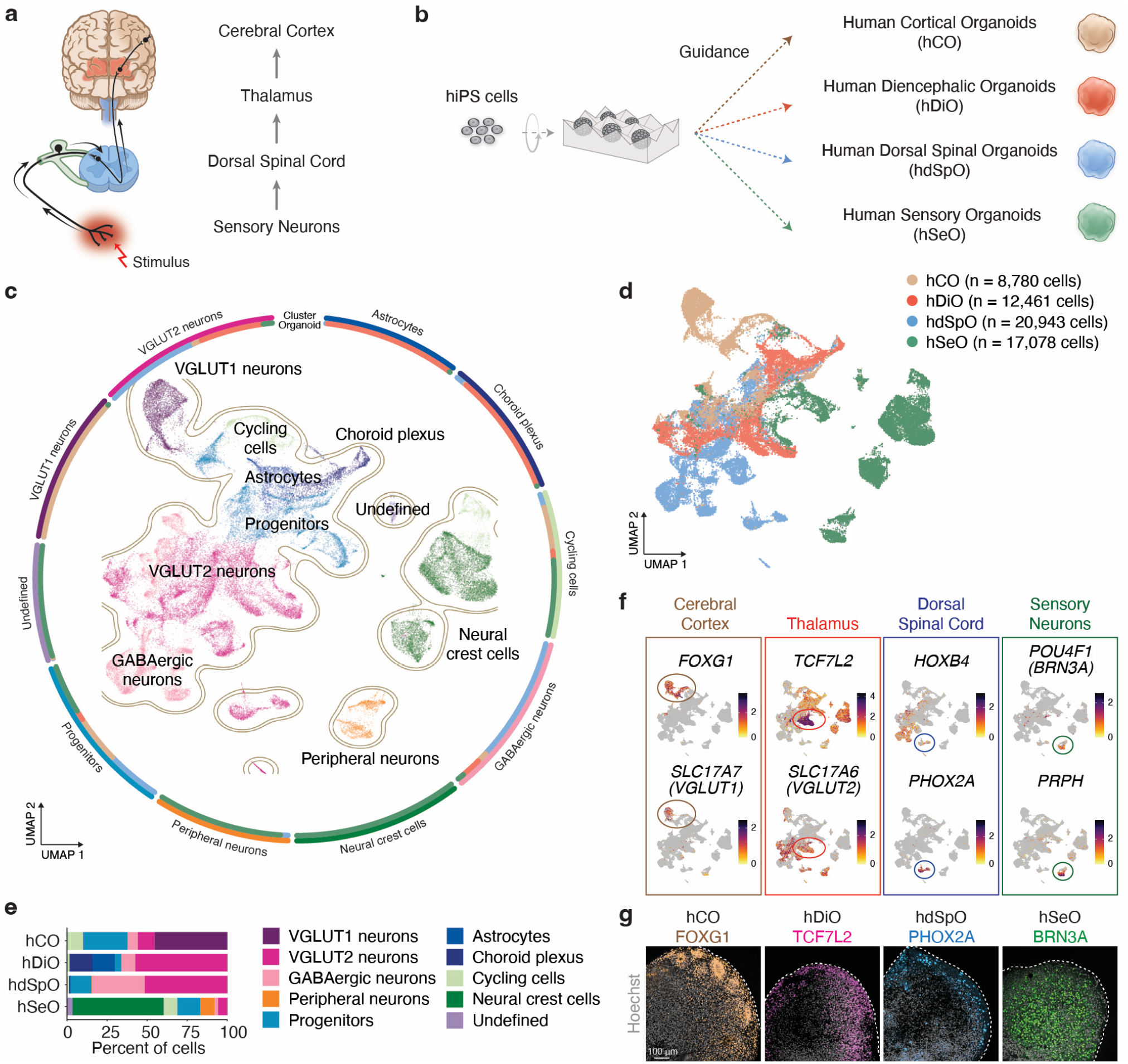
Generation from hiPS cells of components of human ascending sensory pathway. **a**. Schematic illustrating the key components of the ascending anterolateral sensory pathway. **b**. Schematic illustrating the generation of human cortical organoids (hCO), human diencephalic organoids (hDiO), human dorsal spinal cord organoids (hdSpO), and human sensory organoids (hSeO) from hiPS cells. **c**. UMAP visualization of single cell gene expression of hCO, hDiO, hdSpO, and hSeO (hCO, day 83, n = 8,780 cells from 2 hiPS cell lines^34^; hDiO, day 100, n = 12,461 cells from 3 hiPS cell lines^11^; hdSpO, day 68-72, n = 20,943 cells from 5 hiPS cell lines; hSeO; day 68-72, n = 17,078 cells from 4 hiPS cell lines). **d**. UMAP plot colored by the organoid identity. **e**. Cell composition in each regionalized neural organoid based on scRNAseq data. **f**. Gene expression level for regional identity markers. **g**. Immunostaining for specific regional markers in hCO, hDiO, hdSpO, and hSeO at day 63.

## Results

### Generation of the key components of the ascending human spinothalamic pathway

To derive the key parts of the ascending sensory pathway *in vitro*, we generated regionalized neural organoids from human induced pluripotent stem (hiPS) cells using guided differentiation with small molecules and growth factors (Fig. 1b). We first generated human cortical organoids (hCO) and human diencephalic organoids (hDiO) using methods we have previously described and validated^10,11^ (Supplementary Data Fig. 1). To generate human dorsal (posterior) spinal cord organoids (hdSpO), where the soma of spinothalamic projection neurons resides, we modified a protocol we have previously reported for ventral spinal cord organoids^8^ by excluding ventralizing cues (Supplementary Data Fig. 1). Lastly, to derive human somatosensory (also referred here as sensory) organoids (hSeO), we developed a protocol that leveraged cues identified in 2D-based methods^12^ (Supplementary Data Fig. 1). To verify that we have all key cellular components in the spinothalamic tract, we integrated single-cell RNA sequencing data (scRNA-seq) of hCO, hDiO, hdSpO and hSeO (Fig. 1c-e). We found distinct clusters for the key cell types in the ascending circuit: cortical glutamatergic neurons (*FOXG1*^*+*^, *SLC17A7*^*+*^), thalamic excitatory neurons (*TCF7L2*^*+*^, *SLC17A6*^*+*^), dorsal spinothalamic projection neurons (*HOXB4*^*+*^, *PHOX2A*^*+*^), and primary afferent somatosensory neurons (*POU4F1*^*+*^, *PRPH*^*+*^) (Fig. 1f). Lastly, we used immunostaining to confirm protein expression for key region-specific markers in regionalized organoids (Fig. 1g).

### Characterization of human sensory organoids and human dorsal spinal cord organoids

To further characterize the cellular identity of hSeO, we inspected the pattern of expression of several canonical markers using scRNA-seq data (Fig. 2a). We found that approximately 30–40% of neurons (*STMN2*^*+*^) expressed the peripheral neuron-specific markers *POU4F1* and *SIX1*, which we independently confirmed in other batches of organoids by RT-qPCR (Supplementary Data Fig. 2a). We also found a substantial proportion of cells expressing neural crest cell markers, such as *SOX10* and *FOXD3*, which is consistent with a neural crest origin of peripheral sensory neurons^13^ (Fig. 2b, c and Supplementary Data Fig. 2b, c).

**Fig. 2.**
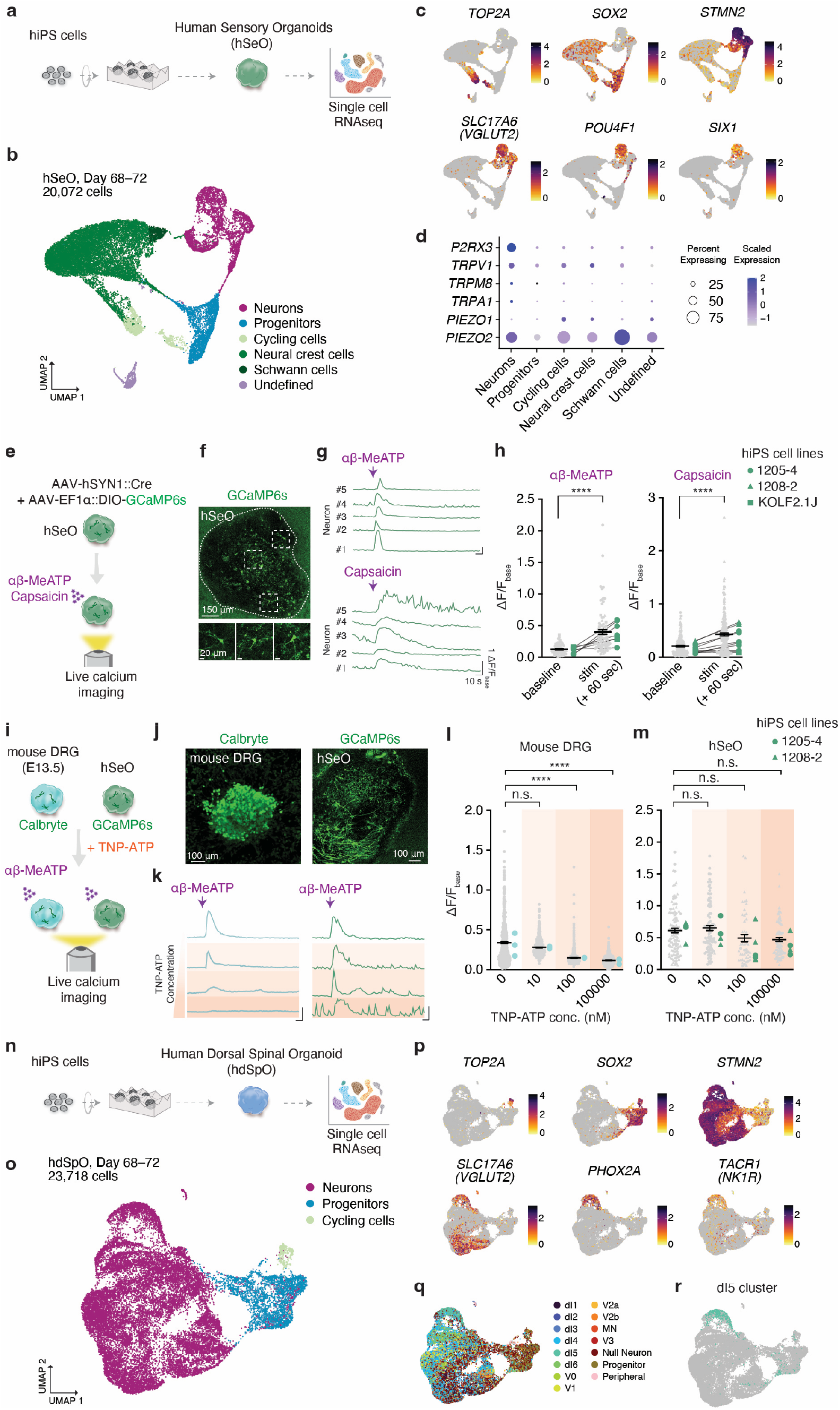
Characterization of hSeO and hdSpO. **a**. Schematic describing the generation of hSeO from hiPS cells and scRNAseq analysis. **b**. UMAP visualization of single cell gene expression of hSeO at day 68–72 (n = 20,072 cells from 4 hiPS cell lines). **c**. UMAP plots showing gene expression of selected cluster-specific markers. **d**. Dot plots showing gene expression of key receptors for sensory stimuli (ATP, *P2RX3*^14^; heat/cold, *TRPV1, TRPM8*^35^; chemical stimulation, *TRPA1*^35^; mechanic stimulation, *PIEZO1, PIEZO2*^36^). **e**. Schematic of live calcium imaging for pharmacological studies. **f**. Representative image showing GCaMP6s expression in hSeO at single cell resolution. **g**. Example ΔF/F_base_ traces before and after agonist treatment showing calcium responses evoked by application of αβ-MeATP (30 μM) and capsaicin (3.3 μM). **h**. Quantification of mean ΔF/F_base_ values (over 60s) at baseline and after exposure to αβ-MeATP (left) (n = 88 neurons from 7 hSeO, 3 hiPS cell lines in 3 differentiations, two-tailed Wilcoxon matched-pairs signed rank test, ****P < 0.0001) and capsaicin (right) (n = 205 cells from 10 hSeO, 3 hiPS cell lines in 3 differentiations, two-tailed Wilcoxon matched-pairs signed rank test, ****P < 0.0001) at day 54–82. **i**. Schematic of live calcium imaging testing various concentrations of TNP-ATP combined with αβ-MeATP exposure in *ex vivo* mouse DRG (E13.5) or hSeO. **j**. Representative images of a Calbryte-incubated mouse DRG (left) and GCaMP6s expression in hSeO (right). **k**. Example ΔF/F_base_ traces demonstrating decrease of αβ-MeATP (30 μM) evoked responses following incubation with TNP-ATP in mouse DRG (left) but not in hSeO (right). The darker orange background represents higher concentration of TNP-ATP. Horizontal scale bar; 20 s. Vertical scale bar; 1 ΔF/F_base_. **l**. Quantification of mean ΔF/F_base_ values for 60 s after exposure to αβ-MeATP (30 μM) in mouse DRG at various concentrations of TNP-ATP (0 nM, n = 518 cells from 3 mouse DRG; 10 nM, n = 605 cells from 3 mouse DRG; 100 nM, n = 652 cells from 3 mouse DRG; 100000 nM, n = 1392 cells from 4 mouse DRG; Kruskal-Wallis test with Dunn’s multiple comparisons test, ****P < 0.0001). **m**. Quantification of mean ΔF/F_base_ values for 60 s after treatment with αβ-MeATP (30 μM) in hSeO at various concentrations of TNP-ATP (0 nM, n = 110 cells from 3 hSeOs; 10 nM, n = 99 cells from 4 hSeOs; 100 nM, n = 50 cells from 6 hSeOs; 100000 nM, n = 77 cells from 4 hSeOs; 2 hiPS cell lines in 2 differentiations at day 75–82; Kruskal-Wallis test with Dunn’s multiple comparisons test). **n**. Schematic showing the generation of hdSpO from hiPS cells and scRNAseq analysis. **o**. UMAP visualization of single cell gene expression of hdSpO at day 68–72 (n = 23,718 cells from 5 hiPS cell lines). **p**. UMAP plots showing gene expression of selected cluster-specific markers. **q**. UMAP plot of hdSpO showing dorsoventral neuron clusters. **r**. UMAP plot of hdSpO showing dI5 cells. Data are shown mean ± s.e.m. Gray dots indicate cells, large colored dots indicate organoids or mouse DRG in g, k, l. Each shape represents a hiPS cell line: Circle, 1205-4; Triangle, 1208-2; Square, KOLF2.1J in h and m.

A unique feature of sensory neurons is their ability to respond to environmental stimuli. To verify whether neurons in hSeO respond to sensory stimuli, we established a live imaging system of intact 3D organoids expressing genetically-encoded calcium indicator using a chamber equipped to inject chemicals. Consistent with high expression of ionotropic calcium–permeable P2X purinoceptor 3, *P2RX3*, and calcium–permeable capsaicin receptor *TRPV1* (Fig. 2d), application of agonists to these receptors, α, β-methyleneATP (αβ-MeATP)^14^ and capsaicin^15^ respectively, induced calcium transients (Fig. 2e–h). Interestingly, the dynamics of the responses were similar between hSeO and intact *ex vivo* primary mouse DRG; αβ-MeATP rapidly induced a fast calcium spike while capsaicin induced a slow and prolonged calcium transient, which is consistent with previous pharmacological studies^13^ (Supplementary Data Fig. 3a–d).

**Fig. 3.**
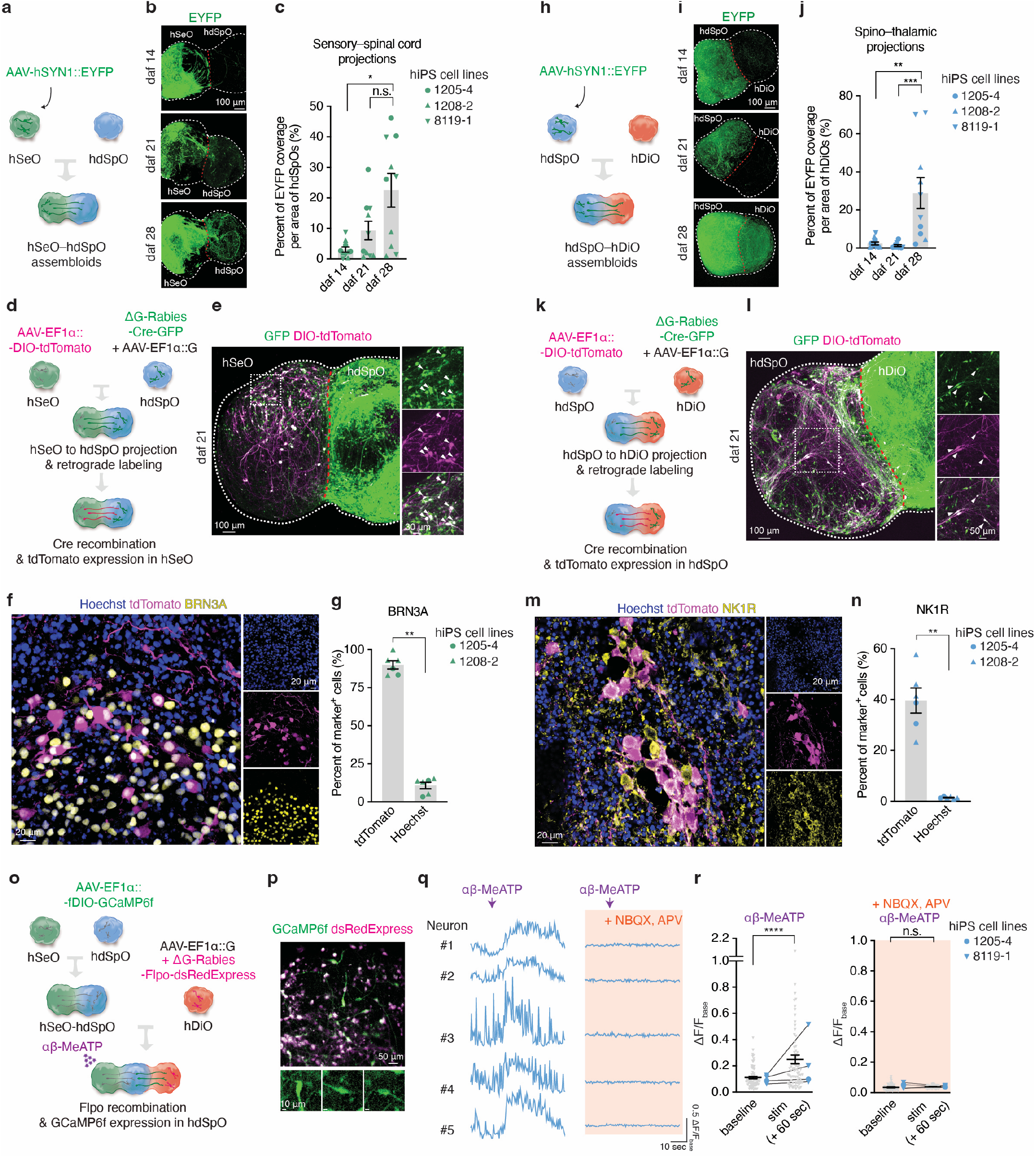
Connectivity in hSeO–hdSpO and hdSpO–hDiO assembloids. **a**. Schematic illustrating the fusion of labeled hSeO and hdSpO to form hSeO–hdSpO assembloids. **b**. Representative images of hSeO–hdSpO assembloids showing a gradual increase in sensory–spinal cord projections over time at daf 14, 21, and 28. **c**. Quantification of EYFP coverage in hdSpO area at daf 14, 21, and 28 in hSeO–hdSpO assembloids (n = 10 assembloids from 3 hiPS cell lines in 3 differentiations; Kruskal-Wallis test with Dunn’s multiple comparisons test, *P = 0.0267). **d**. Schematic illustrating retrograde viral tracing in hSeO–hdSpO assembloid using rabies virus. **e**. Representative image indicating EGFP^+^/tdTomato^+^ cells on the hSeO side of the assembloid at daf 21. **f**. Immunostaining image showing co-localization of BRN3A and tdTomato on the hSeO side of the assembloid. **g**. Percentage of BRN3A^+^ cells out of tdTomato^+^ cells as compared to Hoechst^+^ cells on the hSeO side of the assembloid (n = 6 assembloids from 2 hiPS cell lines in 3 differentiations, two-tailed Mann-Whitney test, **P = 0.0022). **h**. Schematic illustrating the fusion of hdSpO and hDiO to form hdSpO–hDiO assembloids. **i**. Representative images of a hdSpO–hDiO assembloid showing gradual increase of spinothalamic projections at daf 14, 21, and 28. **j**. Quantification of the EYFP in hDiO at daf 14, 21, and 28 in hdSpO–hDiO assembloids (n = 10–11 assembloids from 3 hiPS cell lines in 3 differentiations, Kruskal-Wallis test with Dunn’s multiple comparisons test, **P = 0.0050, ***P = 0.0003). **k**. Schematic illustrating retrograde viral tracing in hdSpO–hDiO assembloid using rabies virus. **l**. Representative image indicating EGFP^+^/tdTomato^+^ cells on the hdSpO side of the assembloid at daf 21. **m**. Immunostaining image showing co-localization of NK1R with tdTomato on the hdSpO side of the assembloid. **n**. Percentage of NK1R^+^ cells out of tdTomato^+^ cells as compared to Hoechst^+^ cells on the hdSpO side of the assembloid (n = 6 assembloids from 2 hiPS cell lines in 3 differentiations, two-tailed Mann-Whitney test, **P = 0.0022). **o**. Schematic illustrating strategy for live calcium imaging of spinothalamic neurons combined with retrograde viral tracing in hSeO–hdSpO–hDiO assembloids. **p**. Representative image of co-expression of GCaMP6f and dsRedExpress on the hdSpO side of hSeO–hdSpO–hDiO assembloids. **q**. Example ΔF/F_base_ traces demonstrating αβ-MeATP (30 μM)-evoked responses (left) and their blockade by exposure to NBQX and APV (50 μM, each) (right). **r**. Quantification of mean ΔF/F_base_ values for baseline and for 60 s after exposure to αβ-MeATP in the absence (left) or presence (right) of NBQX and APV (n = 74 cells from 4 assembloids, 2 hiPS cell lines in 2 differentiations, two-tailed paired t-test, ****P < 0.0001) at daf 70–88. Data are shown mean ± s.e.m. Gray dots indicate cells, large colored dots indicate assembloids in c, g, j, n, r. Each shape represents a hiPS cell line: Circle, 1205-4; Triangle, 1208-2; Inverted triangle, 8119-1 in c, g, j, n, r.

There are species-specific differences in the expression and function of some receptors on sensory DRG neurons, which may be responsible for sensory processing differences and challenges in translation from animal models to humans^7^. For instance, the 2’,3’-O-(2,4,6-trinitrophenyl) adenosine 5’-triphosphate (TNP-ATP) is a potent antagonist of the rodent P2X3 receptor^16^; however, a primate-specific amino acid substitution in the P2X3 receptor is thought to modify sensitivity to TNP-ATP in the macaque DRG^17^. Consistent with previous reports^17^, we found that TNP-ATP successfully blocked calcium responses in ex vivo mouse DRG in a dose-dependent manner. When applied to human sensory neurons in hSeO, however, TNP-ATP had no significant effect (Fig. 2i–m), which is suggestive of differences in responsiveness across species.

We further investigated the cellular identity of neurons in hdSpO using scRNA-seq data (Fig. 2n). Similar to ventralized human spinal cord organoids^8^, hdSpO displayed high neuronal diversity (Fig. 2 o, p and Supplementary Data Fig. 4a, b). Relevant to the ascending somatosensory pathway, we found clusters expressing the spinothalamic projection neuron markers *PHOX2A*^18^ and *TACR1* (encoding NK1R)^19–22^ (Fig. 2p). These cells also express the dI5 marker *LBX1* and *LMX1B* but not the dI4/6 marker *PAX2* (Supplementary Data Fig. 4b). We next annotated neuronal clusters in hdSpO using combinatorial markers expression, as previously described^23,24^ (Fig. 2q,r), and found overlap of *PHOX2A*^*+*^ neurons with dI5^18,25^. Taken together, these transcriptomic and functional studies indicate that these regionalized organoids contain key cellular components of the human ascending sensory pathway.

**Fig. 4.**
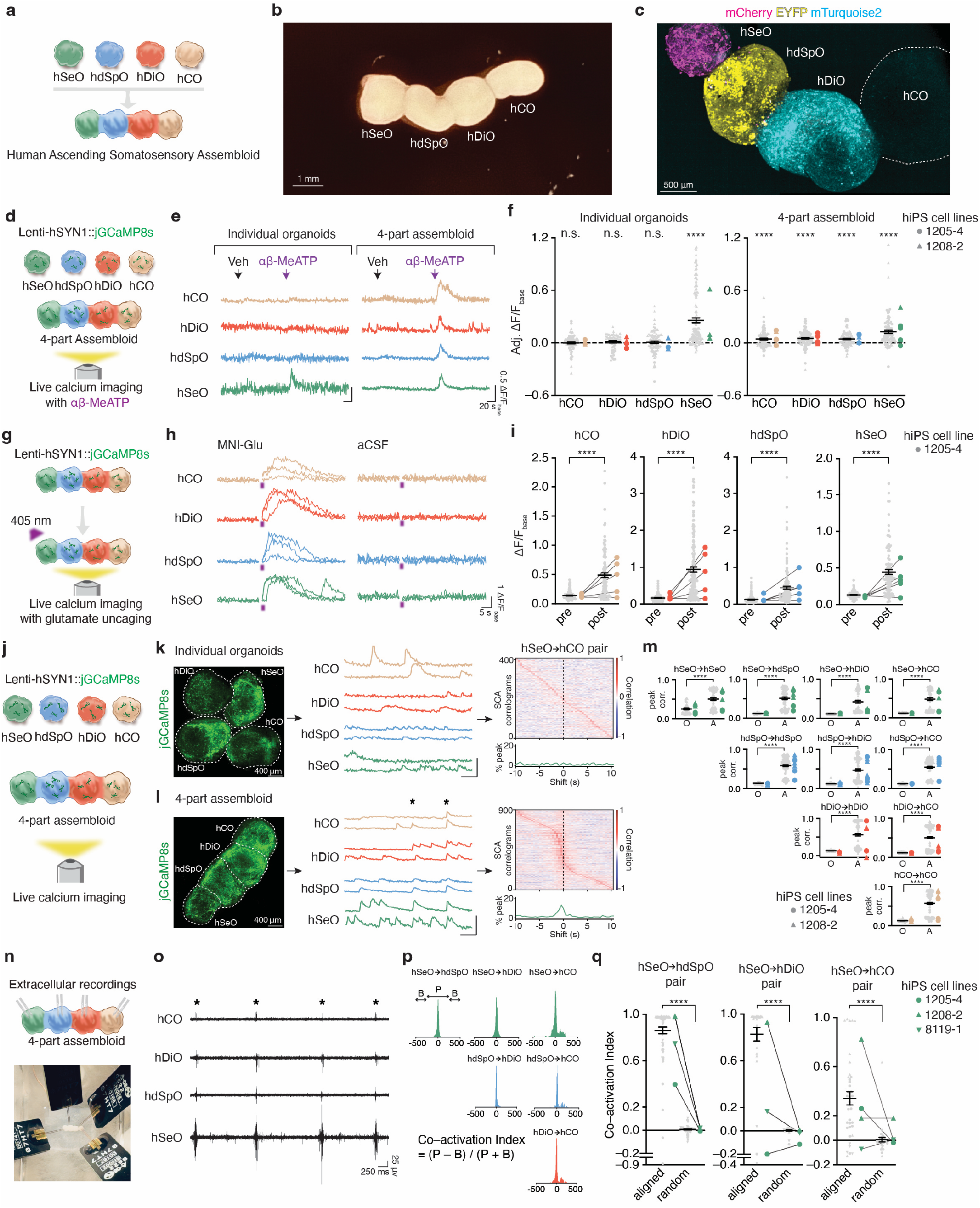
Generation and functional characterization of human ascending sensory assembloids. **a**. Schematic illustrating the fusion of hSeO, hdSpO, hDiO, and hCO to form hASA. **b**. Bright-field image of an hASA at daf 19. **c**. Representative image of an hASA with fluorescent labeling under hSYN1 promoter (hSeO; mCherry, hdSpO; EYFP, hDiO; mTurquoise2). **d**. Schematic illustrating live calcium imaging of individual organoids and four-part assembloid after αβ-MeATP exposure. **e**. Representative ΔF/F_base_ traces demonstrating αβ-MeATP (3 μM)-evoked calcium responses in hSeO in organoids (left). αβ-MeATP (3 μM) induced calcium responses in all regions of assembloids (right). **f**. Quantification of adj.ΔF/F_base_ values as an indicator of calcium responses following αβ-MeATP (3 μM) exposure, compared with null hypothetical value, 0, in individual organoids (left) (hCO, n = 83; hDiO, n = 52; hdSpO, n = 84; hSeO, n = 131 cells from 3–4 organoids, 2 hiPS cell lines in 2 differentiations, two-tailed Wilcoxon signed-rank test, ****P < 0.0001) and assembloids (right) (hCO, n = 91; hDiO, n = 116; hdSpO, n = 98; hSeO, n = 104 cells, from 6 organoids, 2 hiPS cell lines in one differentiation, two-tailed Wilcoxon signed-rank test, ****P < 0.0001) at daf 74–78. **g**. Schematic illustrating glutamate uncaging applied in hASA. UV light (405 nm, 1.31 s) was applied on hSeO cells. **h**. Example ΔF/F_base_ traces from 3 trials showing evoked calcium levels in all four regions with, but not without, MNI-caged glutamate (2.5 mM). **i**. Quantification of mean ΔF/F_base_ values before and after UV illumination (hCO, n = 113; hDiO, n = 157; hdSpO, n = 159; hSeO, n = 109 cells, 5 assembloids, one hiPS cell line in one differentiation, two-tailed Wilcoxon matched-pairs signed rank test, ****P < 0.0001) at daf 74–99. **j**. Schematic illustrating spontaneous live calcium imaging of individual organoids and four-part assembloids. **k**. Individual organoids expressing jGCaMP8s (left), example ΔF/F_base_ traces of cells (middle), and representative SCA correlograms between cells from hSeO and hCO with percentage of peak time-shifts (right). Horizontal scale bar; 10 s, Vertical scale bar; 1 ΔF/F_base_. **l**. Four-part assembloid expressing jGCaMP8s (left), example ΔF/F_base_ traces of cells (middle), and representative SCA correlograms between cells from hSeO and hCO with percentage of peak time-shifts showing a clear peak distribution around 0 (right). Asterisks indicate synchronized calcium events. Horizontal scale bar; 10 sec. Vertical scale bar; 1 ΔF/F_base_. **m**. Correlation values averaged per cell, for combinations of the four regions of hASA. O; organoids, (hCO, n = 83; hDiO, n = 47; hdSpO, n = 54; hSeO, n = 40 cells from 5 non-assembled organoids pairs, 2 hiPS cell lines in one differentiation, two-tailed Mann-Whitney test, ****P < 0.0001), A; assembloids (hCO, n = 176; hDiO, n = 123; hdSpO, n = 111; hSeO, n = 63 cells from 7 assembloids, 2 hiPS cell lines in one differentiation, two-tailed Mann-Whitney test, ****P < 0.0001) at daf 74–99. **n**. Schematic illustrating (upper) and representative picture (bottom) of simultaneous four-part extracellular recordings in an hASA. **o**. Representative traces of spontaneous activity. Asterisks denote synchronous activity across all four regions. **p**. Representative cross-correlograms. Activity counts around peak area (P) and activity counts in base area (B) were used to calculate co–activation index. **q**. Co–activation index values for hSeO→hdSpO, hSeO→hDiO, and hSeO→hCO pairs. Co–activation index values aligned to hSeO activities were significantly higher than values from randomly selected time points (hSeO→hdSpO, n = 88; hSeO→hDiO, n = 34; hSeO→hCO, n = 37 channels, from 3–4 assembloids, 3 hiPS cell lines in 2 differentiations, two-tailed Wilcoxon matched-pairs signed rank test, ****P < 0.0001) at daf 80–83. Data are shown mean ± s.e.m. Gray dots indicate cells in f, i, m or channels in q, large colored dots indicate assembloids in f, i, m, q. Each shape represents a hiPS cell line: Circle, 1205-4; Triangle, 1208-2; Inverted triangle: 8119-1 in f, i, m, q.

### Projections in hSeO–hdSpO and hdSpO–hDiO assembloids

We next investigated the specificity of projections between pairs of regionalized organoids. First, we investigated whether the sensory neurons in hSeO could send projections to spinal neurons in hSeO–hdSpO assembloids. We virally labeled hSeO with hSYN1::EYFP and integrated them with hdSpO to form hSeO–hdSpO assembloids (Fig. 3a). We found that the area of EYFP fluorescence on the hdSpO side, corresponding to projections from hSeO neurons, increased progressively over 28 days after fusion (daf) (Fig. 3b,c). We next used rabies virus-mediated retrograde tracing combined with Cre recombination in hSeO–hdSpO assembloids (Fig. 3d). At daf 21, we observed tdTomato^+^/GFP^+^ cells on the hSeO side, indicating successful retrograde tracing of rabies-Cre-GFP resulting in the recombination and expression of DIO-tdTomato (Fig. 3e). To investigate the specificity of these projections, we co-immunostained for the sensory neuronal marker BRN3A and found that nearly 80% of tdTomato^+^ cells were also BRN3A^+^ (Fig. 3f,g). Notably, when we generated assembloids by integrating hSeO with hCO, we observed only minimal projections from hSeO that did not increase over time (Supplementary Data Fig. 5a–c).

**Fig. 5.**
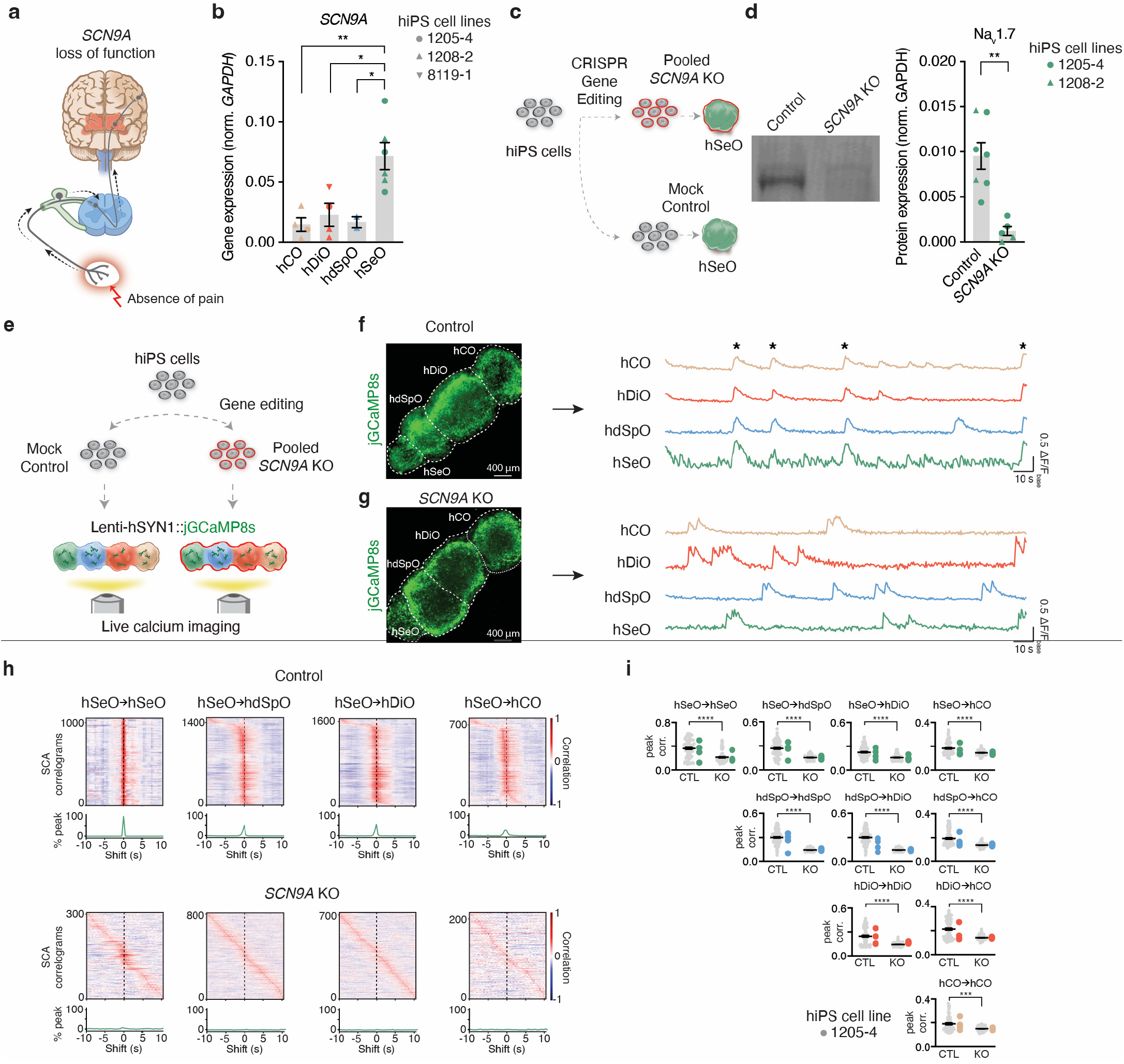
Abnormal calcium activity in SCN9A knock-out hASAs. **a**. *SCN9A* gene loss results in the absence of pain in patients. **b**. RT-qPCR analysis of *SCN9A* expression across four regions (hCO, n = 4; hDiO, n = 4; hdSpO, n = 2; hSeO, n = 6 samples, from 1–3 hiPS cell lines in 2–4 differentiations, one-way ANOVA (F_3, 12_ = 8.153) with Tukey’s multiple comparisons test, *P = 0.0333 for hdSpO vs. hSeO, *P = 0.0164 for hDiO vs. hSeO, **P = 0.0059) at day 45–100. **c**. Schematic illustrating the generation of *SCN9A* KO hiPS cells using CRISPR/Cas9 genome editing and their differentiation into hSeO. **d**. Representative western blotting image and quantification showing Na_v_1.7 expression in control and *SCN9A* KO hSeO at day 48–69 (Control, n = 7; KO, n = 5 samples from 2 hiPS cell lines in 2 differentiations, two-tailed unpaired t-test, **P = 0.0011). **e**. Schematic illustrating derivation and live calcium imaging of control and *SCN9A* KO hASA. **f**. Control assembloids expressing jGCaMP8s (left) and example ΔF/F_base_ traces of cells (right). **g**. *SCN9A* KO assembloids expressing jGCaMP8s (left) and example ΔF/F_base_ traces of cells (right). **h**. Representative SCA correlograms between cells from hSeO→hSeO, hSeO→hdSpO, hSeO→hDiO, and hSeO→hCO for a control assembloid (top) and in a *SCN9A* KO assembloid (bottom). The histograms of peak time-shifts (below each correlograms panel) show peaks around 0 in the control, but not in the *SCN9A* KO assembloid. **i**. Correlation values averaged per cell, for all combinations across four regions. CTL; control assembloids, (hCO, n = 60; hDiO, n = 89; hdSpO, n = 105; hSeO, n = 88 cells from 4 assembloids, one hiPS cell line in one differentiation), KO; *SCN9A* KO assembloids (hCO, n = 100; hDiO, n = 117; hdSpO, n = 131; hSeO, n = 101 cells from 6 assembloids, one hiPS cell line in one differentiation, two-tailed Mann-Whitney test, ***P = 0.0005, ****P < 0.0001) at daf 74–78. Data are shown mean ± s.e.m. Gray dots indicate cells in i, large colored dots indicate organoids, western blot samples, or assembloids in b, d, i. Each shape represents a hiPS cell line: Circle, 1205-4; Triangle, 1208-2; Inverted triangle, 8119-1 in b, d, i.

We took a similar approach to investigate spinothalamic projections in hdSpO–hDiO assembloids (Fig. 3h). We observed a significant increase of spinothalamic projections at daf 28 (Fig. 3i,j). Using rabies virus-mediated retrograde tracing (Fig. 3k,l), we found that approximately 40% of tdTomato^+^ cells co-expressed the spinal cord projection marker NK1R^+^ (Fig. 3m,n). We also observed co-localization of tdTomato and PHOX2A, which is a marker of spinothalamic projection neurons (Supplementary Data Fig. 5d). These studies, in combination with our previous experiments on thalamocortical projections in hCO–hDiO assembloids^11^, confirmed that the three pairs of assembloids built from the four regionalized neural organoids intrinsically recapitulate some of the basic inter-regional projections.

We next investigated the functionality of these connections in three-part hSeO–hdSpO–hDiO assembloids by combining rabies virus-mediated retrograde tracing, recombinase Flpo-dependent expression of GCaMP6f, and live calcium imaging following αβ-MeATP treatment (Fig. 3o). We assembled AAV-EF1α::fDIO-GCaMP6f-infected hdSpO with hSeO to generate hSeO–hdSpO. At daf 43, we infected hDiO with rabies-Flpo-dsRedExpress and AAV-G and fused these onto the hdSpO side of the hSeO–hdSpO assembloid. At daf 70, we detected expression of fDIO-GCaMP6f, which is suggestive of transmission of rabies-Flpo-dsRedExpress from hDiO into hdSpO (Fig. 3p). These GCaMP6f^+^ neurons demonstrated increased calcium activity after αβ-MeATP exposure, thereby indicating signal transmission from sensory neurons to spinal neurons (Fig. 3q,r). These calcium transients were blocked by the glutamate receptor antagonists NBQX and APV (Fig. 3q,r). These results imply that chemical stimulation of sensory neurons in hSeO elicited calcium activity that was transmitted to, and induced calcium increase in connected rabies-Flpo-labeld spinothalamic projection neurons in hdSpO. Moreover, hdSpO neurons projected axons into hDiO, as evidenced by rabies tracing.

### Functional assembly of four-part human ascending sensory assembloids

We generated human ascending sensory assembloids (hASA) by linearly placing the four key neural regionalized organoids in a tilted cell culture plate (Fig. 4a, b). We performed viral labeling of each organoid with distinct hSYN1 promoter-driven fluorescent proteins prior to fusion and confirmed maintenance of distinct domains of these assembloids (Fig. 4c). In one continuous imaging session, we serially treated hASA with either vehicle or αβ-MeATP while recording jGCaMP8s signal at single cell resolution (Fig. 4d). We calculated the adjusted ΔF/F_base_ by subtracting the response to vehicle from the response to αβ-MeATP. We found that αβ-MeATP exposure significantly elevated calcium levels in hASA neurons in all 4 regions at daf 74–78 (Fig. 4e,f and Supplementary Video 1). In contrast, in non-assembled organoids, only neurons had a significant calcium rise after application of αβ-MeATP, while neurons in other organoids did not (Fig. 4e,f). To further verify whether specific stimulation of sensory neurons in hSeO elicited responses in other regions of the assembloid, we used 405 nm photo-stimulation in the presence of MNI-caged glutamate to trigger rapid and localized release of glutamate in a specific area (Fig. 4g). We performed photo-stimulation of hSeO within hASA and detected a robust increase in calcium levels, not only in hSeO but also in all other regions, at daf 74– 99 (Fig. 4h,i). These experiments suggest transmission of somatosensory signals within the assembloid from sensory neurons through spinal and thalamic neurons to cortical neurons.

We next explored whether the four-part assembly is associated with emergent network properties. We recorded baseline calcium activity at single cell resolution and compared hASA with four non-assembled organoids placed in close proximity just before imaging (Fig. 4j-l). Remarkably, we found that neurons in assembloids exhibited synchronous activity across all four regions (Fig. 4l and Supplementary Video 2). We applied scaled correlation analysis (SCA) to minimize effects of low frequency noise signals^26^. Representative SCA correlograms from all possible pairs between hSeO and hCO neurons in an assembloid showed that the correlation peaks were distributed around 0 sec shift, while representative SCA correlograms from unfused organoids had equal distribution of correlation peaks (Fig. 4k,l). Peak correlation for all the combinations of the four regions were significantly higher in assembloids than in individual, non-assembled neural organoids. (Fig. 4m).

We further investigated activity patterns through concomitant extracellular recordings with 32 electrodes in each of the four hASA parts (Fig. 4n). We found spontaneous and synchronous activity across entire assembloid (Fig. 4o). We calculated co–activation indexes based on cross-correlograms^27^ (Fig. 4p). When we compared the co–activation index aligned with activity of hSeO, we found significantly higher values than alignment to random timepoints (Fig. 4q). Taken together, these results demonstrate that the assembly of intact 3D human regionalized neural organoids resembling the human DRG, dorsal spinal cord, thalamus, and cerebral cortex enables the formation of connections capable of synchronized activity across the pathway and in response to somatosensory like (chemical sensory) stimuli.

### Loss of *SCN9A* is associated with abnormal activity in human ascending sensory assembloids

Lastly, we studied whether hASA could be used to identify functional defects in models of neurologic and psychiatric disorders. Pathogenic variants in the gene *SCN9A*, encoding the voltage-gated sodium channel Na_V_1.7, cause pain insensitivity if the variant leads to loss of channel function, or hyperalgesia if the variant causes gain of function^6^ (Fig. 5a). While hypo-excitability of DRG neurons has been proposed as a disease mechanism in *SCN9A* knock-out (KO) cellular and animal models^28–31^, the circuit-level consequences have not yet been explored in a human model.

We first verified expression of *SCN9A* and found high levels in hSeO (Fig. 5b). We then applied CRISPR-mediated gene editing to induce one or more frame-shift variants in the *SCN9A* gene (Fig. 5c). We confirmed that hSeO derived from pooled *SCN9A* KO iPS cell lines exhibited decreased *SCN9A* mRNA and Na_v_1.7 protein levels (Fig. 5d, Supplementary Data Fig. 6a). The expression of the sensory neuron marker BRN3A was not affected by *SCN9A* KO (Supplementary Data Fig. 6b-d). Consistent with previous reports of hypo-excitability of sensory neurons in the absence of this key sodium channel^28^, we found that *SCN9A* KO hSeO displayed reduced frequency of spontaneous calcium activity (Supplementary Data Fig. 6e,f). We then generated hASA from isogenic control and *SCN9A* KO hiPS cells in order to explore the circuit level consequences. We recorded spontaneous activity patterns using jGCaMP8s in all regions from control and *SCN9A* KO hASA (Fig. 5e). We discovered that the synchronous pattern of activity across hASA was reduced in *SCN9A* KO, although non-synchronized events were found in all regions (Fig. 5f,g and Supplementary Video 3, 4). We then performed SCA analysis and found a significant reduction in correlation, indicating that *SCN9A* KO impaired the emergent synchronization across the assembloid (Fig. 5h,i). These results highlight the capability of hASA to capture circuit level dysfunction associated with *SCN9A* KO and thus to model aberrations in emergent properties within human sensory circuits.

## Discussion

Reconstructing the ascending sensory pathway and circuits *ex vivo* holds significant promise for understanding the mechanisms involved in human sensory system development and thereby unraveling the pathophysiology of sensory-related disorders. We previously demonstrated the potential of assembloids composed of regionalized neural organoids derived from hiPS cells to model circuits *in vitro*^8,32,33^. Here, we leveraged this assembloid approach to generate a four-component ascending sensory circuit going from peripheral sensory neurons to cortical neurons. This human ascending sensory assembloid system is the first *in vitro* model encompassing the major components of the human anterolateral spinothalamic pathway, including peripheral sensory neurons, dorsal spinal cord projection neurons, thalamic neurons, and glutamatergic cortical neurons. This allowed us to visualize transmission of chemically triggered calcium signals through the network following sensory stimulus application and to capture emergent properties of the sensory pathway that cannot be seen in individual organoids, such as synchronous activity across the four parts of the assembloid. Importantly, hASA can also capture humanspecific aspects of the sensory system, such as reduced responsiveness to TNP-ATP. hASA therefore hold great promise for addressing disease pathophysiology and testing therapeutic approaches. We discovered that *SCN9A* loss resulted in reduced synchronous activity across the pathway, which suggests that inefficient transmission from sensory neurons may ultimately lead to analgesia in patients. Going forward, this multi-cellular platform can be applied to model a wide range of disorders, including sensory dysfunction in neurodevelopmental disorders or hyperalgesia, and could be helpful for drug screening.

Improvements at several levels would enhance the utility of hASA. First, hiPS cell-derived sensory neurons are still limited in receptor expression and thus the repertoire of responses to sensory stimulation. Future studies could improve the specification and maturation of DRG neurons. Second, there are several neural pathways involved in sensory perception in addition to the anterolateral spinothalamic tract, which our current hASA models. We envision that these other pathways can be built in parallel with hASA or within the same system. Further studies should be conducted to investigate in detail the specificity of connectivity in these assembloids, including within individual organoids, and the role of activity-dependent processes in establishing and refining these neural networks. These efforts could ultimately be used to dissect the intrinsic rules for establishing connectivity in early human circuits, as well as suggest strategies for accelerating maturation in these *in vitro* models.

Overall, this new platform holds potential to yield insights into how the human sensory pathways assemble and therefore to develop and screen therapeutics for pain and other sensory system-related disorders.

## Acknowledgements

We thank members of the Pasca laboratory at Stanford University for scientific input. This work was supported by the Stanford Brain Organogenesis Big Idea Grant from the Wu Tsai Neurosciences Institute (to S.P.P.), the NYSCF Robertson Stem Cell Investigator Award (to S.P.P.), the Kwan Research Fund (to S.P.P.), the Senkut Research Funds (to S.P.P.), the Chan Zuckerberg Initiative Ben Barres Investigator Award (to S.P.P.), the CZ BioHub Investigator Program (to S.P.P.), the Nurturing Next-generation Researchers Program from the National Research Foundation of Korea (to J.K.), NINDS K08-NS123544-01 (to N.D.A), the Brain and Behavior Research Foundation (to N.D.A) and the Foundations of the National Institutes of Health Deeda Blair Research Initiative (to N.D.A.).

## Author contributions

J.K., K.I. and S.P.P. conceived the project and designed experiments. J.K. carried out the differentiation, characterization of regionalized neural organoids, extracellular recordings and calcium imaging. K.I. performed neural differentiations, single cell transcriptomics, calcium imaging, and *SCN9A* gene editing. O.J. contributed to the analysis pipelines for synchronized activities. M.V.T. performed neural differentiations, western blotting, and characterization of regionalized neural organoids. Z.H. analyzed calcium imaging results. N.D.A. contributed to primary tissue experiments.

G.S. contributed to analyze the data. J.K., K.I. and S.P.P. wrote the manuscript with input from all the authors.

## Materials and Methods

### Characterization and maintenance of hiPS cells

Human induced pluripotent stem (hiPS) cell lines used in this study were validated using methods described in previous studies. Genome-wide SNP genotyping was performed using the Illumina genome-wide GSAMD-24v2-0 SNP microarray at the Children’s Hospital of Philadelphia. Cultures were tested for and maintained Mycoplasma free. A total of 5 control hiPS cell lines derived from fibroblasts collected from 5 healthy subjects were used for experiments (Supplementary Table 1). Two of the control hiPS cell lines were used to generate *SCN9A* knock-out hiPS cell lines. Approval for experiments was obtained from the Stanford IRB panel and informed consent was obtained from all subjects.

### Generation of organoids (hCO, hDiO, hdSpO, and hSeO) from hiPS cells

For neural differentiation, hiPS cells were cultured on vitronectin-coated plates (Life Technologies, A14700) in Essential 8 medium (Life Technologies, A1517001). Cells were passaged every 4–5 days with UltraPure™ 0.5 mM EDTA, pH 8.0 (Thermo Fisher Scientific, 15575). For the generation of regionalized neural organoids, hiPS cells were incubated with Accutase (Innovative Cell Technologies, AT104) at 37°C for 7 min and dissociated into single cells. Optionally, 1–2 days before organoid formation, hiPS cells were exposed to 1% dimethylsulfoxide (DMSO) (Sigma-Aldrich, 472301) in Essential 8 medium. For aggregation into organoids, about 3 x 10^6^ single hiPS cells were seeded per AggreWell-800 plate well in Essential 8 medium supplemented with the ROCK inhibitor Y27632 (10 μM, Selleckchem, S1049), centrifuged at 100 *g* for 3 min, and then incubated at 37°C in 5% CO_2_. On the next day (day 1), organoids consisting of approximately 10,000 cells were collected and transferred into ultra-low attachment plastic dishes (Corning, 3262) in Essential 6 medium (Thermo Fisher Scientific, A1516401) supplemented with patterning molecules as shown in Supplementary Data Fig. 1.

hCO were generated as previously described^10,37^. For day 1 to 6, Essential 6 medium was changed every day and supplemented with dorsomorphin (2.5 μM, Sigma-Aldrich, P5499) and SB-431542 (10 μM, R&D Systems, 1614), and was changed every day. On day 7, organoids were transferred to neural medium containing Neurobasal™-A Medium (Thermo Fisher Scientific, 10888022), B-27™ Supplement, minus vitamin A (Thermo Fisher Scientific, 12587010), GlutaMAX™ Supplement (1:100, Thermo Fisher Scientific, 35050079), Penicillin-Streptomycin (1:100, Thermo Fisher Scientific, 15070063), supplemented with FGF2 (20 ng/ml, R&D Systems, 233-FB) and EGF (20 ng/ml, R&D Systems, 236-EG) until day 22. From day 23 to day 46, the neural media was supplemented with BDNF (20 ng/ml, PeproTech, 450-02), NT3 (20 ng/ml, PeproTech, 450-03), L-Ascorbic Acid 2-phosphate

Trisodium Salt (AA; 200 μM, Wako, 323-44822), N6, 2’-O-Dibutyryladenosine 3’,5’-cyclic monophosphate sodium salt (cAMP; 50 μM, Millipore Sigma, D0627), and cis-4, 7, 10, 13, 16, 19-Docosahexaenoic acid (DHA; 10 μM, Millipore Sigma, D2534). From day 47, neural medium containing Neurobasal™-A Medium was used for media changes (every 4 days).

hDiO were generated as previously described^11^. For day 1 to 6, Essential 6 medium was changed every day and supplemented with dorsomorphin and SB-431542. On day 5, medium was additionally supplemented with 1 μM CHIR (Selleckchem, S1263). On day 7, organoids were maintained in neural media supplemented with CHIR. On day 9 of differentiation, neural medium was supplemented with 100 nM SAG (day 9–15, Millipore Sigma, 566660-1MG), in addition to the CHIR. Furthermore, on day 12–18 of differentiation, organoids were supplemented with 30 ng/mL BMP7 (PeproTech, 120-03P), in addition to the compounds described above. From day 19 to 46, the neural medium was supplemented with BDNF, NT3, AA, cAMP, and DHA. For day 19–25, DAPT (2.5 μM, STEMCELL Technologies, 72082) was additionally added. From day 47, neural medium containing Neuro-basal™-A Medium was used for media changes (every 4 days).

To generate hdSpO, from day 1 to day 6, organoids were maintained Essential 6 medium supplemented with SB-431542 and dorsomorphin. From day 5 to day 6, CHIR (3 μM) was added. On day 7, organoids were transferred to a neural medium supplemented with EGF, Retinoic acid (0.1 μM, Sigma-Aldrich, R2625), and CHIR (3 μM, Selleckchem, S1263). On day 22, the media was supplemented with BDNF, cAMP, AA, IGF-1 (10 ng/ml, PeproTech, 100-11) until the end of experiments. From day 22 to day 28, DAPT was added.

To generate hSeO, from day 1 to day 6 in suspension, organoids were maintained in Essential 6 medium, supplemented with SB-431542. From day 1 to day 4, BMP4 (5 ng/ml, PeproTech, 120-05ET) was added. From day 3 to day 5, CHIR (600 nM) was added. On day 6 in suspension, organoids were transferred to neural medium supplemented with SB-431542. From day 9, BDNF, GDNF (25 ng/ml PeproTech, 450-10), and NGF (25 ng/ml, PeproTech, 450-01) were supplemented until the end of the experiment. From day 9 to day 14, medium was supplemented with SB-431542. From day 15 to day 20, DAPT was included.

### Cryosection and immunocytochemistry

Organoids and assembloids were fixed in 4% paraformaldehyde (PFA)/phosphate buffered saline (PBS) overnight at 4°C. They were then washed in PBS and transferred to 30% sucrose/PBS for 2–3 days until the organoids/assembloids sank into the solution. Subsequently, they were rinsed in optimal cutting temperature (OCT) compound (Tissue-Tek OCT Compound 4583, Sakura Finetek) and 30% sucrose/PBS (1:1), embedded, and snap-frozen using dry ice. For immunofluorescence staining, 30 μmthick sections were cut using a Leica Cryostat (Leica, CM1860). Cryosections were washed with PBS to remove excess OCT from the sections and blocked in 10% Normal Donkey Serum (NDS, Millipore Sigma, S30-100ML), 0.3% Triton X-100 (Millipore Sigma, T9284-100ML), and 1% BSA diluted in PBS for 1 hour at room temperature. The sections were then incubated overnight at 4°C with primary antibodies diluted in PBS containing 2% NDS and 0.1% Triton X-100. PBS was used to wash the primary antibodies and the cryosections were incubated with secondary antibodies in PBS with the PBS containing 2% NDS and 0.1% Triton X-100 for 1 hour. The following primary antibodies were used for staining: anti-FOXG1 (rabbit, Takara, M227, 1:400 dilution), anti-TCF7L2 (rabbit, Cell Signaling Technology, 2569S, 1:200 dilution), anti-PHOX2A (rabbit, Abcam, ab155084, 1:200 dilution), anti-BRN3A (mouse, Sigma, MAB1585, 1:200 dilution), anti-mCherry (goat, Biorbyt, orb153320, 1:1000 dilution), anti-NK1R (rabbit, Sigma, S8305, 1:200 dilution). Alexa Fluor dyes (Life Technologies) were used at 1:200 to 1:1,000 dilution and nuclei were visualized with the Hoechst 33258 dye (Life Technologies, H3549, 1:10,000 dilution). Cryosections were mounted for microscopy on glass slides using Aqua-Poly/Mount (Polysciences, 18606) or Vectashield (Vector labs, H-1000), and imaged on a confocal microscope. Images were processed and analyzed using Fiji (ver. 2.14.0) and IMARIS (Oxford Instruments).

### Real-time qPCR

Three to five organoids were collected in the same tube and considered as one sample. RNA from samples was isolated using the RNeasy Plus Mini kit (Qiagen, 74136). Template cDNA was prepared by reverse transcription using the SuperScriptTM III First-Strand Synthesis SuperMix for qRT-PCR (Thermo Fisher Scientific, 11752250). qPCR was performed using the SYBRTM Green PCR Master Mix (Thermo Fisher Scientific, 4312704) on a QuantStudio™ 6 Flex Real-Time PCR System (Thermo Fisher Scientific, 4485689). Primers used in this study are listed in Supplementary Table 2.

### Single cell RNA-seq library preparation and data analysis

Dissociation of organoids was performed as described previously^10,33,38,39^. For the organoid dissociation, 4–5 randomly selected organoids were pooled to obtain single cell suspension and then incubated in 30 U/mL papain enzyme solution (Worthington Biochemical, LS003126) and 0.4% DNase (12,500 U/mL, Worthington Biochemical, LS2007) at 37°C for 45 min. After enzymatic dissociation, organoids were washed with a solution including protease inhibitor and gently triturated to achieve a single cell suspension. Cells were resuspended in 0.04% BSA/PBS (Millipore-Sigma, B6917-25MG) and filtered through a 70 μm Flowmi Cell Strainer (Bel-Art, H13680-0070) and the number of cells were counted. To target 7,000 cells after recovery, approximately 11,600 were loaded per lane on a Chromium Single Cell 3′chip (Chromium Next GEM Chip G Single Cell Kit, 10x Genomics, PN-1000127) and cDNA libraries were generated with the Chromium Next GEM Single Cell 3′ GEM, Library & Gel Bead Kit v3.1 (10x Genomics, PN-1000128 and PN-1000269), according to the manufacturer’s instructions. Each library was sequenced using the Illumina NovaSeq S4 2 × 150 bp by Admera Health. UMI counting was performed by the ‘count’ function (--include-introns=TRUE) in Cell Ranger (v7.0.1 and 7.1.0) with GRCh38/Ensembl 98 reference. Further downstream analyses were performed using the R package Seurat (v4.3.0). Genes on the X or Y chromosome were removed from the count matrix to avoid biases in clustering due to the sex of the hiPS cell lines. Cells with more than 10,000 or less than 2,500 detected genes, with less than 2,000 UMI counts, or cells with mitochondrial content higher than 15% were excluded. Genes that were not expressed in at least three cells were not included in the analysis. Gene expression was normalized using a global-scaling normalization method (normalization method, ‘LogNormalize’; scale factor, 10,000), and the 2,000 most variable genes were selected (selection method, ‘vst’) and scaled (mean = 0 and variance = 1 for each gene). hSeO samples (n = 4) and hdSpO samples (n = 5) were each separately integrated using ‘FindIntegrationAnchors’ and ‘IntegrateData’ functions with the default parameter. The top 10 principal components were used for clustering (resolution of 1.0), using the ‘FindNeighbors’ and ‘FindClusters’ functions, and for visualization with UMAP. Clusters were grouped based on the expression of known marker genes.

For comparison across four region organoids, hSeO samples (n = 4) and hdSpO samples (n = 5) from this study, along with hDiO samples^11^ (n = 3) and hCO samples^34^ (n = 2), were aggregated using the ‘aggr’ function (--normalize=mapped) in Cell Ranger. Cells with more than 7,500 or less than 2,000 detected genes, with less than 2,000 UMI counts, or with mitochondrial content higher than 15% were excluded. Normalization, variable gene detection, and scaling were performed as described above. The top 50 principal components were used for clustering (resolution of 1.0), using the ‘FindNeighbors’ and ‘FindClusters’ functions, and for visualization with UMAP. A circular plot was generated using the R package plot1cell^40^.

The classification of dorsoventral neuronal cell types of hdSpO was performed based on the combinatorial expression of known markers as previously described^23,24^ by using ‘doCellPartition’ function (cell_level_min_step1 = 2, cell_level_min_step2 = 1; available at github.com/juliendelile/MouseSpinalCordAtlas).

### Data and materials availability

Data is available at GEO accession number GSE251892.

### Generation of assembloids

Regionalized organoids that were at least day 47 were used to make assembloids. For two-part assembloids, organoids of interest were fused by placing them in proximity in 1.5 ml Eppendorf tubes for 2–3 days in an incubator. For three-or four-part assembloids, the organoids of interest were integrated by placing them in close proximity in tilted 6-well ultra-low attachment plates (Corning, #3471) for 1 week in an incubator. After assembly, the assembloids were maintained in ultra-low attachment plates (24-well or 6-well) with medium change every 4 days. For three-or four-part assembloids, the 6-well ultra-low attachment plates were kept tilted in incubator.

### Viral labeling and rabies tracing

Viral infection of organoids was performed as previously described^41^. Briefly, two or three organoids were placed in a 1.5 ml Eppendorf tube containing 200 μl media with 0.5 μl of the virus and incubated with overnight at 37°C, 5% CO_2_. Next day, 800 μL of fresh culture medium was added. The following day, neural organoids were transferred into fresh culture medium in ultra-low attachment plates. For rabies virus retrograde tracing, organoids representing the ‘presynaptic’ part were labeled with AAV-DIO-tdTomato and organoids representing the ‘postsynaptic’ part were separately labeled with Rabies-ΔG-Cre-GFP and AAV-EF1α::CVS-G. Two days after viral infection, organoids were assembled. After 3 weeks of integration, assembloids were fixed with 4% paraformaldehyde and processed for immunocytochemistry.

For rabies virus tracing and calcium imaging experiments, hdSpO were labeled with AAV-EF1α::fDIO-GCaMP6f, and hDiO were separately labeled with both Rabies-ΔG-Flpo-dsRedExpress and AAV-EF1α::CVS-G. For GCaMP virus infection of four-part assembloids, one single assembloid was cultured overnight in 200 μl medium containing 0.5 μl of the virus of interest, at 37°C, 5% CO_2_. Next day, 4 ml of fresh culture medium was added.

The viruses used in this study were: AAV-DJ-hSYN1::EYFP (Stanford University Neuroscience Gene Vector and Virus Core, GVVC-AAV-16), AAV-DJ-EF1α::CVS-G (Produced by Stanford University Neuroscience Gene Vector and Virus Core using Addgene, plasmid #67528), AAV-DJ-EF1-DIO-tdTomato (Stanford University Neuroscience Gene Vector and Virus Core, GVVC-AAV-169), AAV-DJ-hSYN1::mTurquoise2 (Produced by Stanford University Neuroscience Gene Vector and Virus Core using Addgene, plasmid #99125), AAV-DJ-hSYN1::mCherry (Stanford University Neuroscience Gene Vector and Virus Core, GVVC-AAV-17), Rabies-ΔG-Cre-GFP (Salk), Rabies-ΔG-Flpo-dsRedExpress (Salk, using Addgene, plasdmid #32650), AAV1-EF1α::fDIO-GCaMP6f (Addgene, #128315-AAV1), Lenti-hSYN1::jGCaMP8s (Produced by Vectorbuilder), AAV1-hSYN1::Cre (Addgene, #105553), AAV-DJ-EF1α::DIO-GCaMP6s (Stanford University Neuroscience Gene Vector and Virus Core, GVVC-AAV-91).

### Live calcium imaging from organoids, assembloids

To achieve single-cell resolution live calcium imaging in intact 3D organoids or assembloids without significant sample movement during chemical injection, we acutely attached samples to Ethyleneimine polymer (PEI)-coated coverslips before imaging. Autoclaved 12 mm coverslips were coated with 0.01875% PEI (Sigma, #03880) in water for at least 1 hour at 37°C and then washed three times with water before use. An organoid or assembloid was carefully positioned on top of a PEI-coated coverslip, and an L-shaped tubing-type Chamlide CMB chamber (Live Cell Instrument, CM-B12-1PB) was assembled. This chamber was connected to a peristaltic pump (Harvard Apparatus, #702027) to establish continuous suction through the outlet hole. To prevent sample drying, a vehicle solution was applied immediately after assembly. Organoids and assembloids with detectable expression of genetically-encoded calcium indicators and displaying baseline activity were imaged.

The live imaging chamber was transferred to the confocal microscope stage (Leica Stellaris 5 or SP8), and the samples were imaged under controlled environmental conditions (37°C, 5% CO_2_) using a 5x or 10x objective. We waited for 5–10 minutes, and then started time-lapse live imaging at approximately 0.436 seconds per frame, with adjustments for each sample. For imaging spontaneous activity patterns, calcium activity was recorded for 3–5 minutes. To capture calcium responses after chemical injection, we recorded 60s baseline activity. Following this, αβ-MeATP (Tocris, #3209) or capsaicin (Tocris, #0462) dissolved in vehicle was injected over 30 s at a slow speed (approximately 2 ml/min) to minimize any potential movement of the samples.

For analysis, regions of interest (ROIs) were manually drawn, and raw fluorescent intensities were exported using Fiji (ver. 2.14.0). Using MATLAB (ver. R2023a), raw fluorescent intensities were transformed into relative changes in fluorescence: ΔF/F_base_ = (F(t) - F_base_) / F_base_, where F_base_ is the lower 5^th^ percentile value of the session. The mean amplitudes of ΔF/F_base_ from each cell for 60 s were compared before and after chemical injection.

For Fig. 2g,h, the neural media supplemented with BDNF (20 ng/mL), GDNF, and NGF (25 ng/ml each) was used as vehicle solution. After 1 min of baseline imaging, 30 μM of αβ-MeATP or 3 μM of capsaicin was injected into the chamber.

For Fig. 2k-l, the neural media supplemented with BDNF (20 ng/mL), GDNF, and NGF (25 ng/ml each) was used as vehicle solution. hSeO were incubated with various concentrations of TNP-ATP (10, 100, or 100000 nM) dissolved in the vehicle solution. After 1 min of baseline imaging, 30 μM of αβ-MeATP was injected into the chamber.

For Fig. 3q,r, assembloids were acutely attached to a PEI-coated 12 well cell culture plate before the imaging. The plate was moved onto the microscope stage, and the imaging was done as the lid of the plate was open. Neural media supplemented with BDNF, GDNF, and NGF was used as vehicle solution. After 1 min baseline imaging in 1 ml of vehicle solution, 500 μl of αβ-MeATP (90 μM) was added during imaging to make 30 μ? of αβ-MeATP finally. After 6-min-long imaging, samples were incubated with NBQX (50 μM, Tocris, #0373) and APV (50 μM, Tocris, #0106) for 30 min in the incubator. Calcium responses by αβ-MeATP in the presence of NBQX and APV was performed as described above.

For Fig. 4e,f, DPBS with calcium, magnesium (Cytiva, SH30264.01) or artificial cerebrospinal fluid (aCSF) containing 124 mM NaCl, 3 mM KCl, 1.25 mM NaH_2_PO_4_, 1.2 mM MgSO_4_, 1.5 mM CaCl_2_, 26 mM NaHCO_3_ and 10 mM D-(+)-glucose with addition of GlutaMAX (Gibco, 35050061) was used as a vehicle solution. After 1 min baseline imaging, 3 μM of αβ-MeATP was injected into the chamber.

For Fig. 4k,l, aCSF was used as a vehicle solution.

For Fig. 5f,g, aCSF was used as a vehicle solution.

To compute correlation with SCA, signals were partitioned in chunks of 10s. Then, the correlation between co-occurring signal chunks was evaluated using Pearson’s r. Finally, the overall correlation between the signals was given by the average of the r values across all signal chunks. This approach yields an accurate correlation estimate between signals on a short timescale, irrespective of common activations over longer periods of time.

### Glutamate uncaging in assembloids

?ssembloids were imaged under environmentally controlled conditions (37°C) using a 5x objective in a confocal microscope (Leica SP8). For glutamate uncaging experiments, MNI-caged-L-glutamate (Tocris, 1490) was used at a final concentration of 2.5 mM in aCSF. The FRAP software module of the Leica SP8 confocal microscope was used to uncage glutamate using UV light (405 nm). At a frame rate of 2.3 frames/sec, one trial consisted of 50 frames for pre-stimulation, 3 frames of UV stimulation (in specified ROI) and 100 frames for post-stimulation, and 3 trials were applied for one sample. Calcium responses were calculated comparing mean of ΔF/F_base_ during 50 frames (21.8 seconds) before and after UV illumination as described above, and averaged values of 3 trials were used for each sample.

### Western blotting

Three to five organoids were collected in the same tube and considered as one sample. Whole cell protein lysates for samples were prepared using RIPA buffer system (Santa Cruz, sc-24948). Protein concentration was quantified using the bicinchoninic acid assay (Pierce, Thermo Fisher, catalogue no. 23225). For electrophoresis, 30 μg of protein per lane was loaded and run on a 4–12% Bis-Tris PAGE gel (Bolt 4–12% Bis-Tris Protein Gel, Invitrogen, catalogue no. NW04122BOX) and transferred onto a polyvinyl difluoride membrane (Immobilon-FL, EMD Millipore). Membranes were blocked with 5% BSA in tris-buffered saline with Tween (TBST) for 1 h at room temperature and incubated with primary antibodies; anti-GAPDH (glyceraldehyde 3-phosphate dehydrogenase) (mouse, 1:5,000, GeneTex, GTX627408) for 1 day, and anti-SCN9A (rabbit, 1:1,000, Alomone Labs, ASC-008) for 6 days. Membranes were washed three times with TBST and then incubated with near-infrared fluorophore-conjugated species-specific secondary antibodies: Goat Anti-Mouse IgG Polyclonal Antibody (IRDye 680RD, 1:10,000, LI-COR Biosciences, 926–68070) or Goat Anti-Rabbit IgG Polyclonal Antibody (IRDye 800CW, 1:10,000, LI-COR Biosciences, 926– 32211) for 1 h at room temperature. Following secondary antibody application, membranes were washed three times with TBST, once with TBS and then imaged using a LI-COR Odyssey CLx imaging system (LI-COR) with Image studio software (Licor, v.5.2). Western blots were analyzed using the Image studio software (Licor, v.5.2) or Fiji (ver. 2.14.0). See Source Data for full blots.

### Axon projection imaging and analysis

EYFP^+^ projections in target regions were imaged under environmentally controlled conditions in intact assembloids using a Leica TCS SP8 or Leica Stellaris 5 confocal microscope with a motorized stage. Assembloids were transferred to a well in a 24-well glass bottom plate (Cellvis, P24-0-N) in cell culture medium, and incubated in an environmentally controlled chamber for 15–30 min before imaging. Images were taken using a 10x objective to capture the entire target area at a depth of 100 μm. Projections were quantified using Fiji (ver. 2.14.0). ROIs were manually drawn to cover the target area to be measured in max projection confocal stacks. The percentage of fluorescence positive pixels over the target region area was calculated in binary images.

### Mouse primary tissue dissection and imaging

Mouse tissue samples were obtained under a protocol approved by the Research Compliance Office at Stanford University. Pregnant CD1 mice were obtained from Charles River. Dorsal root ganglia (DRG) were dissected from E13.5 embryos, placed *en bloc* onto PEI-coated 12-well plates, and cultured with DRG culture media (neural media supplemented with BDNF, GDNF, NGF, and 10% FBS). Calcium imaging was performed the next day after dissection.

For Supplementary Data Fig. 3a–d, mouse DRG was incubated with Calbryte520AM (10 μM, AAT Bioquest, #20653). After 45 minutes, the medium was changed to 1 ml of DRG culture medium. To image calcium activity of mouse DRG, a whole-cell culture plate containing mouse DRG was moved to the environment controlled confocal microscope stage. Imaging was done as the lid of the plate was open. After 1 min of baseline line imaging, 500 μl of αβ-MeATP (90 μM) or capsaicin (10 μM) dissolved DRG culture media was added into the well by pipetting without touching the cell culture plate (to prevent XYZ movement). The imaging was performed for a total of 6 min. The final concentrations of the agonists were αβ-MeATP (30 μM) and capsaicin (3.3 μM).

For Fig. 2l,m, mouse DRG were incubated with Calbrye520AM as described above, and additionally incubated with various concentrations of TNP-ATP (10, 100, or 100000 nM, Tocris, #2464) dissolved into DRG culture media for 30 min in the incubator. After 1 min of baseline imaging with TNP-ATP, 500 μl of αβ-MeATP (90 μM) in DRG culture medium was added into the well. The imaging was performed for a total of 6 min.

### Extracellular recordings

Assembloids were embedded into 3% low-melting gel agarose (IBI Scientific, IB70056) and transferred to aCSF. Embedded samples were placed on a Brain Slice Chamber-2 (Scientific Systems Design Inc, S-BSC2) and perfused with aCSF (bubbled with 95% O_2_ and 5% CO_2_) at 37°C. Temperature was controlled and retained at 37°C by connecting to a Proportional Temperature Controller PTC03 (AutoMate Scientific, S-PTC03). Acute 32 channel P-1 probes with 2 shanks (Cambridge NeuroTech, ASSY-37 P-1) were connected to an Acute probe adaptor; 32 channels Samtec to Omnetics (Cambridge NeuroTech, ADPT A32-Om32). Baseline activities without any stimulation were acquired using the Intan 1024ch recording controller (Intan Technologies) at 30,000 Hz for 5 min. Raw recording data were processed using Intan Technologies code and analyzed by MATLAB (version R2022a, MathWorks).

To analyze extracellular recording data, bandpass filtering (350 Hz – 2000 Hz) was performed followed by common median referencing. A threshold (6 times multiplied by standard deviation) for each channel was used to count neuronal activities. Activity from hdSpO, hDiO or hCO was counted ± 500 ms of each activity of hSeO channels. (Activity counts around peak area (P) – activity counts in base area (B)) / (P + B) were used to as co– activation index^27^. Parameters (bin size; 5 ms, plot size; ± 500 ms) were used to draw the representative cross–correlograms.

### Genome editing of hiPS cells to generate *SCN9A* knock-out hiPS cell lines

For generating SCN9A knock-out cells, three sgRNAs were designed and synthesized by Synthego to induce one or more fragment deletions. sgRNA sequences are as follows: 5′-AGCUCGUGUAGCCAUAAUCA-3′, 5′-CGUGUGUAGUCAGUGUCCAG-3′, 5′-UUCUCUUGGUACUCAC-CUGU-3′. hiPS cells were dissociated with accutase and 0.5 million cells were mixed with 300 pmol sgRNAs and 40 pmol Cas9 protein (Synthego, SpCas9 2NLS Nuclease (1000 pmol)). Nucleofection was performed using the P3 Primary Cell 4D-Nucleofector™ X Kit S (Lonza, V4XP-3032), a 4D-nucleofector core unit, and the X unit (Lonza) (program CA-137). Cells were then seeded onto vitronectin-coated 6-well plates in Essential 8 medium supplemented with the ROCK inhibitor Y27632 (10 μM). Essential 8 medium was used for daily medium change. The genotype of pooled cells was determined by PCR with the primer set Fw, 5′-CGAGAAC-TACCCATATTATTAGTGATGG-3′; Rv, 5′-CCAAGAAC-TATCACAAAACGTCTGT-3′, and Sanger sequencing (Genewiz) was performed with the reverse primer.

### Statistics

Data are presented as mean ± s.e.m. unless described otherwise. Raw data were tested for normality of distribution and statistical analyses were performed by unpaired t-test, Mann-Whitney test, paired t-test, Wilcoxon matched-pairs signed rank test, Wilcoxon signed-rank test, one-way ANOVA, and Kruskal-Wallis test with multiple comparison tests depending on the dataset. Sample sizes were estimated empirically. GraphPad Prism Version 10.0.0 or MATLAB (version R2023a, MathWorks) were used for statistical analyses.

**Supplementary Data Fig. 1.**
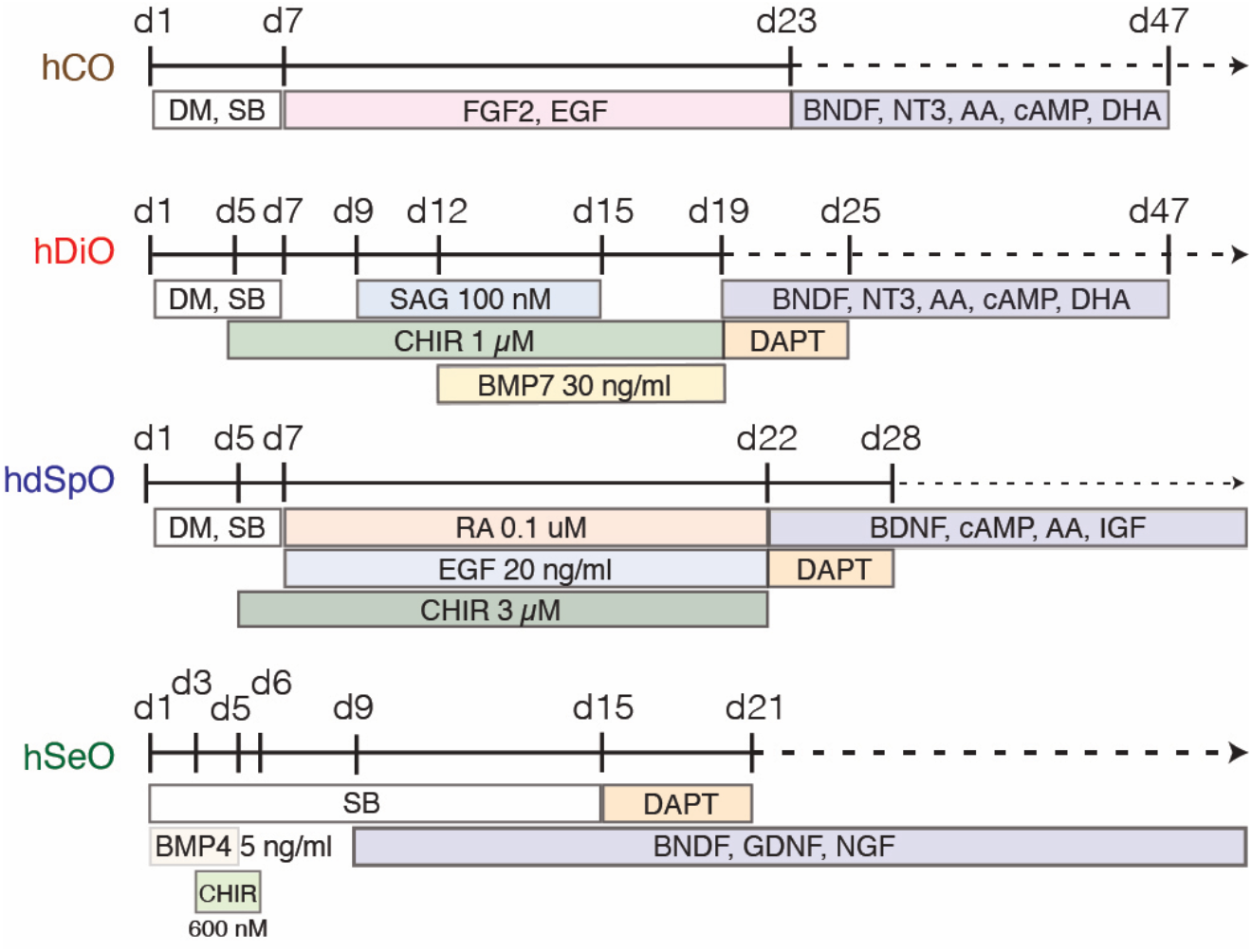
Protocol for generating regionalized neural organoids. Protocols to generate hCO, hDiO, hdSpO, and hSeO from hiPS cells.

**Supplementary Data Fig. 2.**
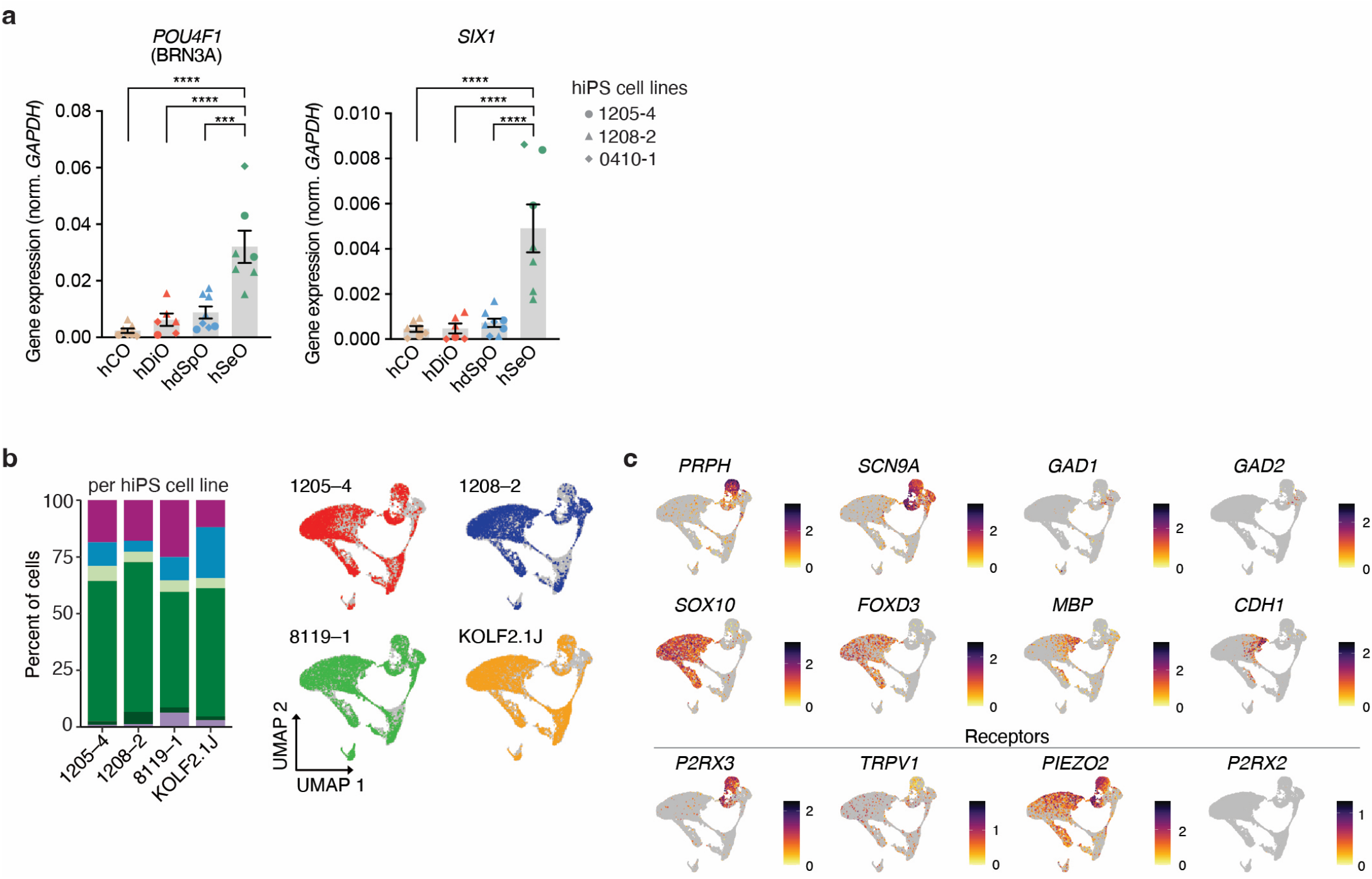
Transcriptomic analysis of hSeO. **a**. RT-qPCR of *POU4F1* (hCO, n = 7; hDiO, n = 6; hdSpO, n = 8; hSeO, n = 7 samples, from 3 hiPS cell lines in 2–4 differentiations, one-way ANOVA, F_3, 24_ = 16.59 with Tukey’s multiple comparisons test, ***P = 0.0001, ****P < 0.0001) and *SIX1* (hCO, n = 7; hDiO, n = 6; hdSpO, n = 8; hSeO, n = 7 samples, from 3 hiPS cell lines in 2–4 differentiations, one-way ANOVA, F_3, 24_ = 15.49 with Tukey’s multiple comparisons test, ****P < 0.0001) expression across four regions at day 45–100. **b**. Cell type composition in hSeO samples colored by cluster (left), and UMAP plots of hSeO samples colored by the hiPS cell line (right). **c**. UMAP plots of hSeO samples showing cell type markers. Data are shown mean ± s.e.m. Each shape represents a hiPS cell line: Circle, 1205-4; Triangle, 1208-2; Diamond, 0410-1 in a.

**Supplementary Data Fig. 3.**
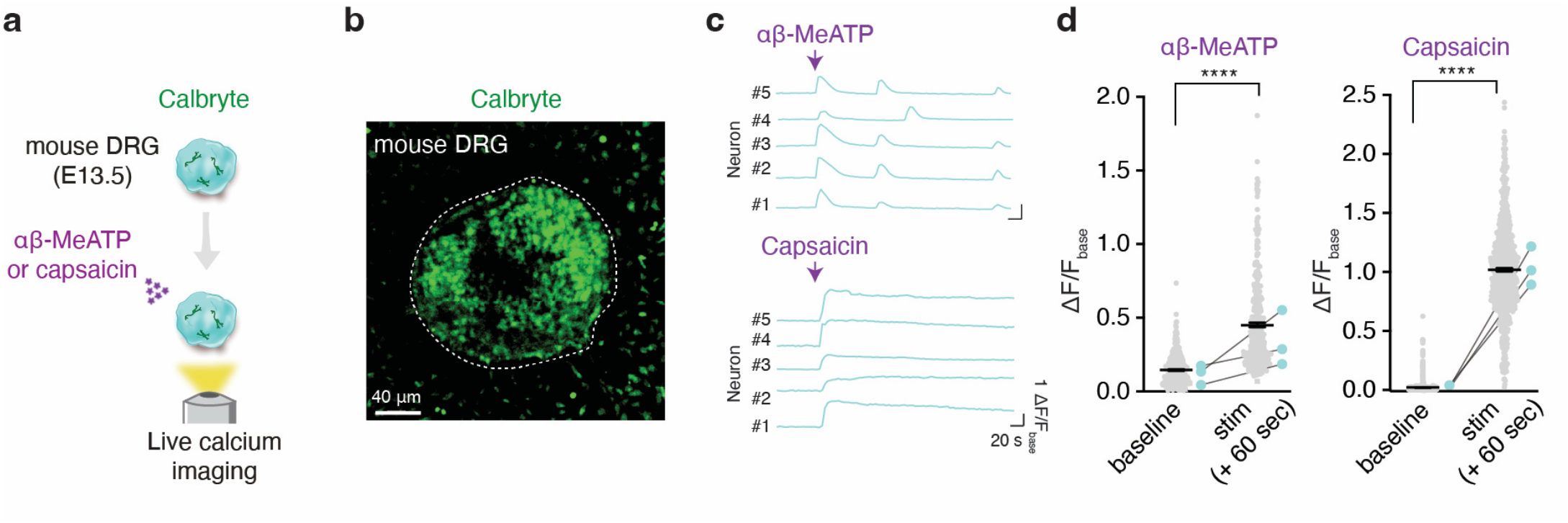
Calcium imaging of isolated mouse DRG. **a**. Schematic of live calcium imaging of isolated mouse DRG following pharmacological treatment. **b**. Representative image showing GCaMP6s expression in mouse DRG. **c**. Example ΔF/F_base_ traces before and after agonist exposure showing calcium responses evoked by αβ-MeATP (30 μM) and capsaicin (3.3 μM). **d**. Quantification of mean ΔF/F_base_ values at baseline and at 60 s after exposure to αβ-MeATP (left) (n = 384 cells from 3 DRG, Two-tailed Wilcoxon matched-pairs signed rank test, ****P < 0.0001) and capsaicin (right) (n = 762 cells from 10 hSeO, 3 DRG, two-tailed Wilcoxon matched-pairs signed rank test, ****P < 0.0001). Data are shown mean ± s.e.m. Gray dots indicate cells, large colored dots indicate DRG in d.

**Supplementary Data Fig. 4.**
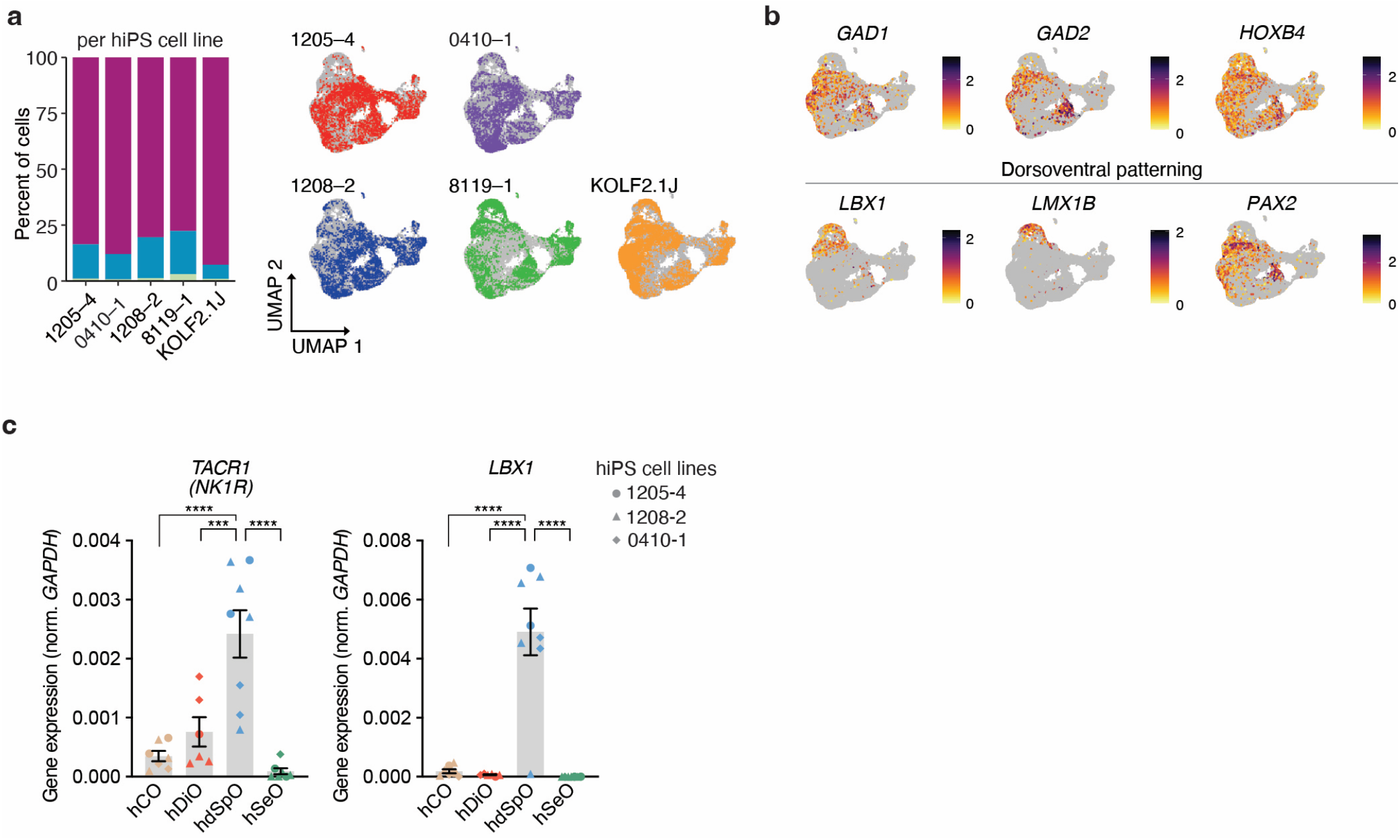
Transcriptomic analysis of hdSpO. **a**. Cell type composition of hdSpO colored by cell clusters (left), and UMAP plots of hdSpO samples colored by the hiPS cell line (right). **b**. UMAP plots of hdSpO samples showing cell type markers. **c**. RT-qPCR of *TACR1* (hCO, n = 7; hDiO, n = 6; hdSpO, n = 8; hSeO, n = 7 samples, from 3 hiPS cell lines in 2–4 differentiations, one-way ANOVA, F_3, 24_ = 17.70 with Tukey’s multiple comparisons test, ***P = 0.0009, ****P < 0.0001) and *LBX1* (hCO, n = 7; hDiO, n = 6; hdSpO, n = 8; hSeO, n = 7 samples, from 3 hiPS cell lines in 2–4 differentiations, one-way ANOVA, F_3, 24_ = 30.50 with Tukey’s multiple comparisons test, ****P < 0.0001) expression across four regions at day 45–100. Data are shown mean ± s.e.m. Each shape represents a hiPS cell line: Circle, 1205-4; Triangle, 1208-2; Diamond, 0410-1 in c.

**Supplementary Data Fig. 5.**
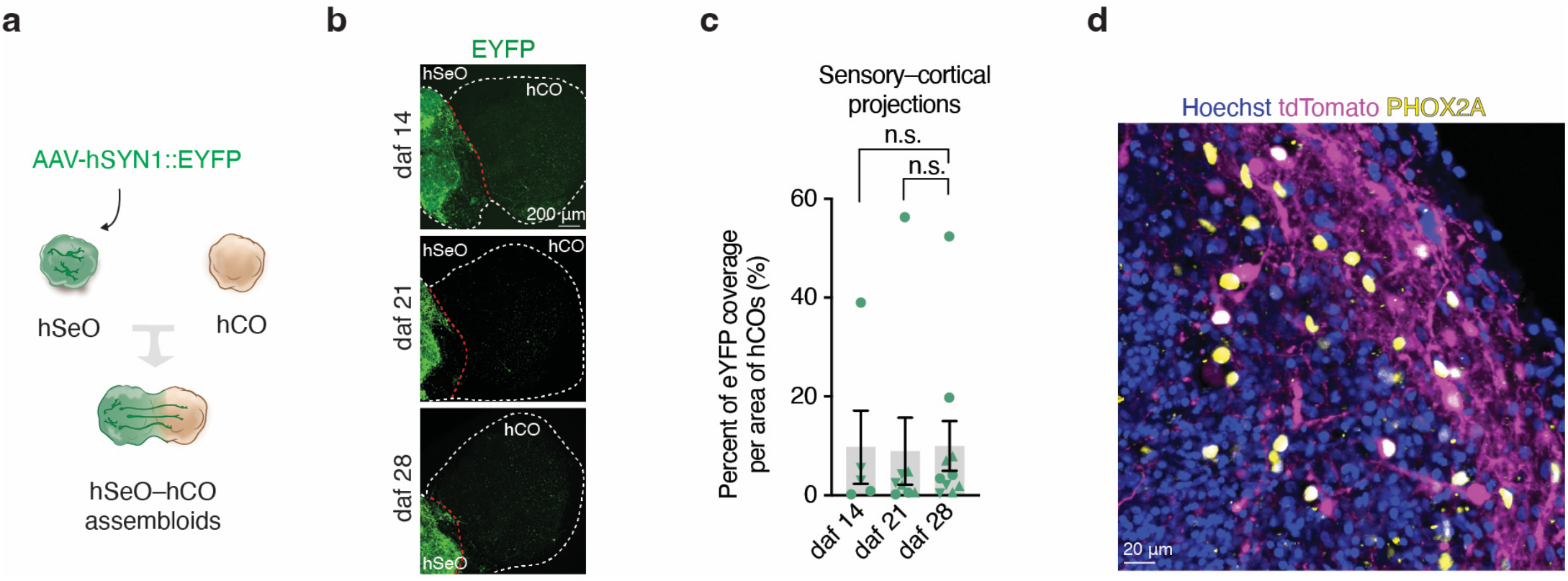
Characterization of projections in assembloids. **a**. Schematic illustrating the fusion of hSeO and hCO to form hSeO-hCO assembloids. **b**. Representative images of a hSeO-hCO assembloid showing a limited number of hSeO-derived hSYN1::EYFP projections at 14, 21, and 28 daf. **c**. Quantification of hSeO-derived EYFP fluorescence in hCO area at 14, 21, and 28 daf in hSeO-hCO assembloids (n = 5–10 assembloids from 3 hiPS cell lines in 3 differentiations, Kruskal-Wallis test with Dunn’s multiple comparisons test, n.s.; not significant). **c**. Immunostaining image showing co-localization of PHOX2A and tdTomato on the hdSpO side of hSeO–hdSpO assembloids. Data are shown mean ± s.e.m. Each shape represents a hiPS cell line: Circle, 1205-4; Triangle,1208-2; Inverted triangle, 8119-1 in c.

**Supplementary Data Fig. 6.**
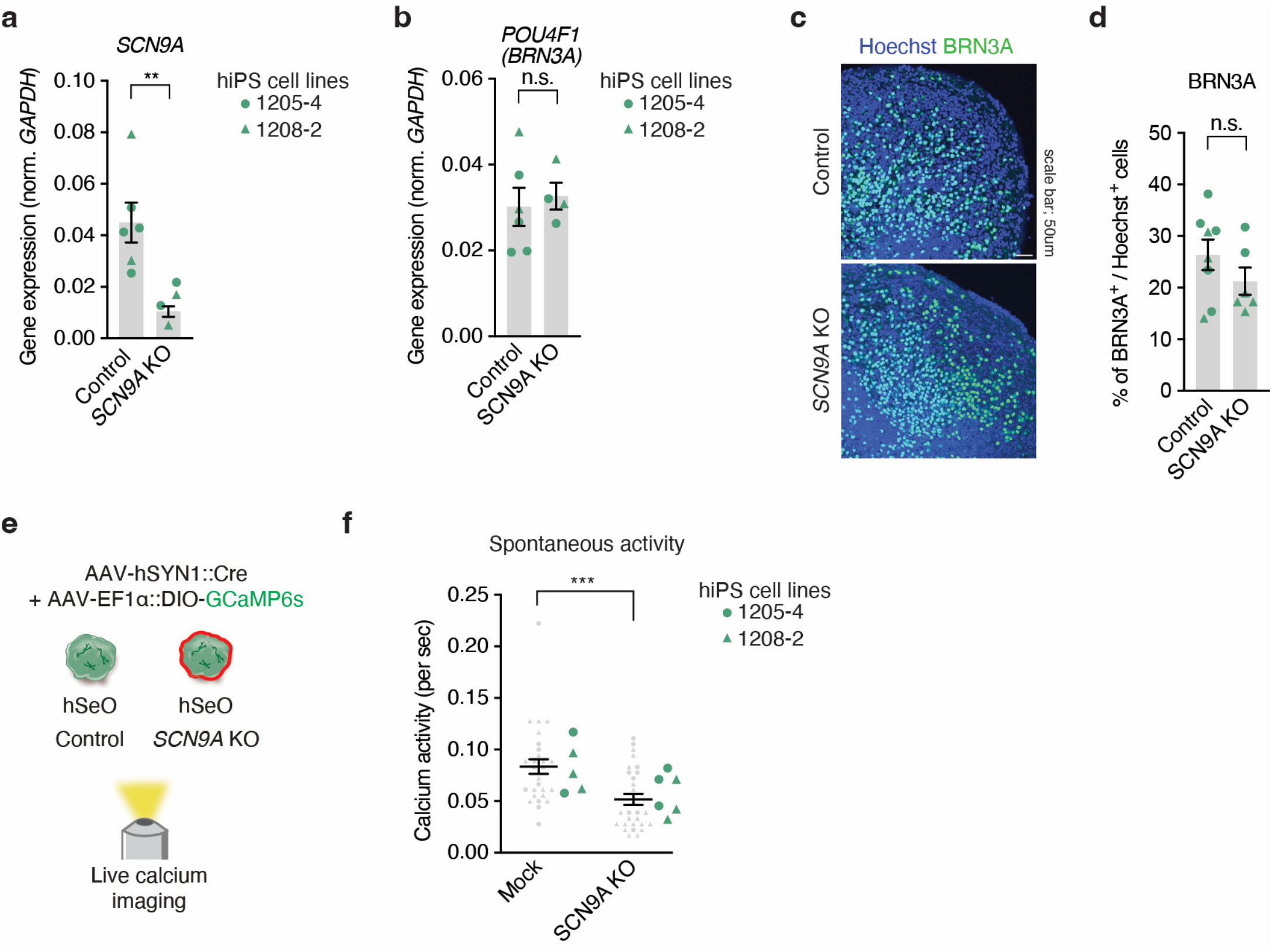
Characterization of SCN9A KO hSeO. **a**. RT-qPCR analysis of *SCN9A* expression in control and *SCN9A* KO hSeO (n = 4–6 samples from 2 hiPS cell lines in 2 differentiations, two-tailed unpaired t-test, **P = 0.0082) at day 48–69. **b**. RT-qPCR analysis of *BRN3A* expression in control and *SCN9A* KO hSeO (n = 4–6 samples from 2 hiPS cell lines in 2 differentiations, two-tailed unpaired t-test, n.s.; not significant) at day 48–69. **c**. Immunocytochemistry images for BRN3A in control and *SCN9A* KO hSeO. **d**. Quantification of the percentage of BRN3A^+^ cells in control and *SCN9A* KO hSeO (n = 6–8 samples from 2 hiPS cell lines in 2 differentiations, two-tailed unpaired t-test, n.s.; not significant). **e**. Calcium imaging of control and *SCN9A* KO hSeO **f**. Quantification of the number of spontaneous calcium responses (CTL, n = 28 cells from 5 organoids; KO, n = 29 cells from 6 organoids, 2 hiPS cell lines in 2 differentiations, two-tailed Mann-Whitney test, ***P = 0.0006) at day 64-82. Data are shown mean ± s.e.m. Gray dots indicate cells, large colored dots indicate organoids in f. Each shape represents a hiPS cell line: Circle, 1205-4; Triangle, 1208-2 in a, b, d, and f.

**SUPPLEMENTARY TABLE 1.** hiPS cell lines used in this study.

| Experiment / Line | 1205-4 | 1208-2 | 8119-1 | KOLF2.1J | 0410-1 |
| --- | --- | --- | --- | --- | --- |
| hSeO single cell RNA seq | O | O | O | O |  |
| hdSpO single cell RNA seq | O | O | O | O | O |
| qPCR | O | O |  | O |  |
| hSeO calcium imaging | O | O |  | O |  |
| Projection imaging | O | O | O |  |  |
| Rabies tracing + Immunostaining | O | O |  |  |  |
| Assembloid calcium imaging | O | O |  |  |  |
| Assembloid extracellular recording | O | O | O |  |  |
| SCN9A gene editing | O | O |  |  |  |

**SUPPLEMENTARY TABLE 2.**
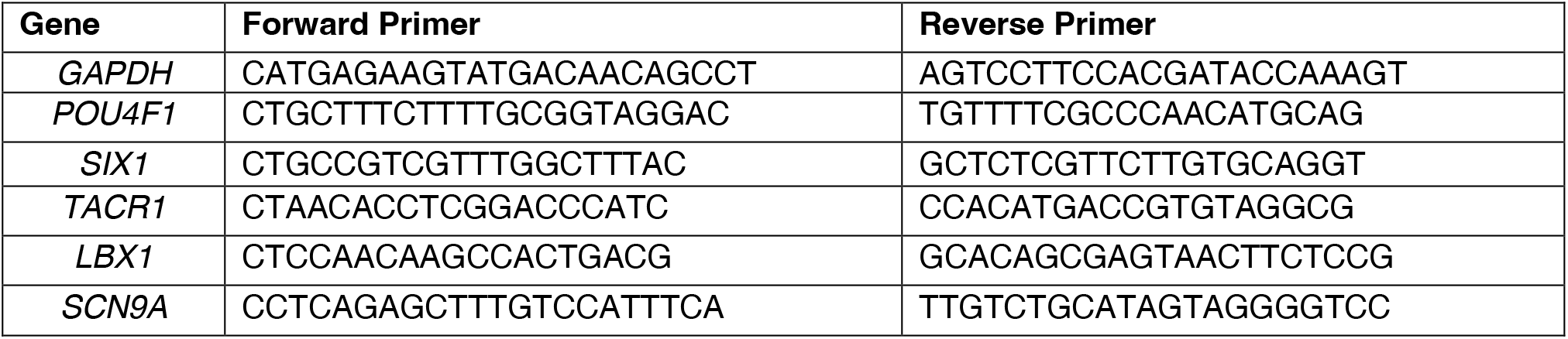
Primers used for qPCR experiments.

## References

1. Colloca, L. et al. Neuropathic pain. Nat Rev Dis Primers 3, 17002 (2017).

2. Basbaum, A. I., Bautista, D. M., Scherrer, G. & Julius, D. Cellular and Molecular Mechanisms of Pain. Cell 139, 267–284 (2009).

3. Han, Q. et al. SHANK3 Deficiency Impairs Heat Hyperalgesia and TRPV1 Signaling in Primary Sensory Neurons. Neuron 92, 1279–1293 (2016).

4. Dawes, J. M. et al. Immune or Genetic-Mediated Disruption of CASPR2 Causes Pain Hypersensitivity Due to Enhanced Primary Afferent Excitability. Neuron 97, 806–822.e10 (2018).

5. Orefice, L. L. et al. Peripheral Mechanosensory Neuron Dysfunction Underlies Tactile and Behavioral Deficits in Mouse Models of ASDs. Cell 166, 299–313 (2016).

6. Drenth, J. P. H. & Waxman, S. G. Mutations in sodium-channel gene SCN9A cause a spectrum of human genetic pain disorders. Journal of Clinical Investigation 117, 3603–3609 (2007).

7. Mogil, J. S. Animal models of pain: progress and challenges. Nat Rev Neurosci 10, 283–94 (2009).

8. Andersen, J. et al. Generation of Functional Human 3D Cortico-Motor Assembloids. Cell 183, 1913–1929.e26 (2020).

9. Kayalioglu, G. Projections from the Spinal Cord to the Brain. in The Spinal Cord 148–167 (2009).

10. Pasca, A. M. et al. Functional cortical neurons and astrocytes from human pluripotent stem cells in 3D culture. Nat Methods 12, 671–678 (2015).

11. Kim, J.-I. et al. Human assembloids reveal the consequences of CACNA1G gene variants in the thalamocortical pathway. bioRxiv (2023) doi:10.1101/2023.03.15.530726.

12. Zimmer, B. et al. Human iPSC-derived trigeminal neurons lack constitutive TLR3-dependent immunity that protects cortical neurons from HSV-1 infection. Proceedings of the National Academy of Sciences 115, (2018).

13. Meltzer, S., Santiago, C., Sharma, N. & Ginty, D. D. The cellular and molecular basis of somatosensory neuron development. Neuron 109, 3736–3757 (2021).

14. North, R. A. The P2X3 subunit: a molecular target in pain therapeutics. Curr Opin Investig Drugs 4, 833–40 (2003).

15. Caterina, M. J. et al. The capsaicin receptor: a heat-activated ion channel in the pain pathway. Nature 389, 816–24 (1997).

16. Virginio, C., Robertson, G., Surprenant, A. & North, R. A. Trinitro-phenyl-substituted nucleotides are potent antagonists selective for P2X1, P2X3, and heteromeric P2X2/3 receptors. Mol Pharmacol 53, 969–73 (1998).

17. Serrano, A. et al. Differential Expression and Pharmacology of Native P2X Receptors in Rat and Primate Sensory Neurons. The Journal of Neuroscience 32, 11890–11896 (2012).

18. Roome, R. B. et al. Phox2a Defines a Developmental Origin of the Anterolateral System in Mice and Humans. Cell Rep 33, 108425 (2020).

19. Choi, S. et al. Parallel ascending spinal pathways for affective touch and pain. Nature 587, 258–263 (2020).

20. Todd, A. J. Neuronal circuitry for pain processing in the dorsal horn. Nat Rev Neurosci 11, 823–836 (2010).

21. Nichols, M. L. et al. Transmission of Chronic Nociception by Spinal Neurons Expressing the Substance P Receptor. Science (1979) 286, 1558–1561 (1999).

22. Mantyh, P. W. et al. Inhibition of Hyperalgesia by Ablation of Lamina I Spinal Neurons Expressing the Substance P Receptor. Science (1979) 278, 275–279 (1997).

23. Rayon, T., Maizels, R. J., Barrington, C. & Briscoe, J. Single-cell transcriptome profiling of the human developing spinal cord reveals a conserved genetic programme with human-specific features. Development 148, (2021).

24. Delile, J. et al. Single cell transcriptomics reveals spatial and temporal dynamics of gene expression in the developing mouse spinal cord. Development (Cambridge) 146, (2019).

25. Qian, Y., Shirasawa, S., Chen, C.-L., Cheng, L. & Ma, Q. Proper development of relay somatic sensory neurons and D2/D4 inter-neurons requires homeobox genes Rnx/Tlx-3 and Tlx-1. Genes Dev 16, 1220–33 (2002).

26. Nikolić, D., Mureşan, R. C., Feng, W. & Singer, W. Scaled correlation analysis: a better way to compute a cross-correlogram. European Journal of Neuroscience 35, 742–762 (2012).

27. Mizuseki, K. & Buzsaki, G. Theta oscillations decrease spike synchrony in the hippocampus and entorhinal cortex. Philos Trans R Soc Lond B Biol Sci 369, 20120530 (2014).

28. McDermott, L. A. et al. Defining the Functional Role of NaV1.7 in Human Nociception. Neuron 101, 905–919.e8 (2019).

29. Minett, M. S. et al. Distinct Nav1.7-dependent pain sensations require different sets of sensory and sympathetic neurons. Nat Commun 3, 791 (2012).

30. Gingras, J. et al. Global Nav1.7 Knockout Mice Recapitulate the Phenotype of Human Congenital Indifference to Pain. PLoS One 9, e105895 (2014).

31. Deng, L. et al. Nav1.7 is essential for nociceptor action potentials in the mouse in a manner independent of endogenous opioids. Neuron 111, 2642–2659.e13 (2023).

32. Miura, Y. et al. Generation of human striatal organoids and cortico-striatal assembloids from human pluripotent stem cells. Nat Bio-technol 38, 1421–1430 (2020).

33. Birey, F. et al. Assembly of functionally integrated human forebrain spheroids. Nature 545, 54–59 (2017).

34. Khan, T. A. et al. Neuronal defects in a human cellular model of 22q11.2 deletion syndrome. Nat Med 26, 1888–1898 (2020).

35. Julius, D. TRP Channels and Pain. Annu Rev Cell Dev Biol 29, 355–384 (2013).

36. Murthy, S. E., Dubin, A. E. & Patapoutian, A. Piezos thrive under pressure: mechanically activated ion channels in health and disease. Nat Rev Mol Cell Biol 18, 771–783 (2017).

37. Yoon, S. J. et al. Reliability of human cortical organoid generation. Nat Methods 16, 75–78 (2019).

38. Sloan, S. A. et al. Human Astrocyte Maturation Captured in 3D Cerebral Cortical Spheroids Derived from Pluripotent Stem Cells. Neuron 95, 779–790.e6 (2017).

39. Sloan, S. A., Andersen, J., Paşca, A. M., Birey, F. & Paşca, S. P. Generation and assembly of human brain region–specific three-di-mensional cultures. Nat Protoc 13, 2062–2085 (2018).

40. Wu, H. et al. Mapping the single-cell transcriptomic response of murine diabetic kidney disease to therapies. Cell Metab 34, 1064–1078.e6 (2022).

41. Miura, Y. et al. Engineering brain assembloids to interrogate human neural circuits. Nat Protoc 17, 15–35 (2022).

